# Comprehensive genotyping of Brazilian Cassava (*Manihot esculenta* Crantz) Germplasm Bank: insights into diversification and domestication

**DOI:** 10.1101/2020.07.13.200816

**Authors:** Alex C. Ogbonna, Luciano Rogerio Braatz de Andrade, Eder Jorge de Oliveira, Lukas A. Mueller, Guillaume J. Bauchet

## Abstract

Cassava (*Manihot esculenta* Crantz) is a major staple root crop of the tropics, originating from the Amazonas region. In this study, 3,354 cassava landraces and modern breeding lines from the Embrapa Cassava Germplasm Bank (CGB) were characterized. All individuals were subjected to genotyping-by-sequencing (GBS), identifying 27,045 Single Nucleotide Polymorphisms (SNPs). Identity-by-state and population structure analyses revealed a unique set of 1,536 individuals and 10 distinct genetic groups with heterogeneous linkage disequilibrium (LD). On this basis, 1,300 to 4,700 SNP markers were selected for large quantitative trait loci (QTL) detection. Identified genetic groups were further characterized for population genetics parameters including minor allele frequency (MAF), observed heterozygosity (*H_o_*), effective population size estimate 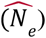 and polymorphism information content (PIC). Selection footprints and introgressions of *M. glaziovii* were detected. Spatial population structure analysis revealed five ancestral populations related to distinct Brazilian ecoregions. Estimation of historical relationships among identified populations suggest earliest population split from Amazonas to Atlantic forest and Caatinga eco-regions and active gene flows. This study provides a thorough genetic characterization of *ex situ* germplasm resources from cassava center of origin, South America, with results shedding light on Brazilian cassava characteristics and its biogeographical landscape. These Findings support and facilitate the use of genetic resources in modern breeding programs including implementation of association mapping and genomic selection strategies.

**Key message:** Brazilian cassava diversity was characterized through population genetics and clustering approaches, highlighting contrasted genetic groups, and spatial genetic differentiation.

## Introduction

Cassava (*Manihot esculenta* ssp. *esculenta*) domestication resulted from human-mediated selection in the Amazonian region and adjacent areas of northern Bolivia (Clement et al. 2010), initially in the early Holocene period (Lombardo et al. 2020), with significant changes in root tuberisation. Evidence suggests that cultivated cassava was domesticated once from *M. esculenta* ssp. *flabellifolia* (K. M. Olsen and Schaal 1999; K. Olsen and Schaal 2001; Kenneth M. Olsen 2004; Schaal, Olsen, and Luiz J C 2006). This hypothesis was further validated by Léotard and colleagues (Léotard et al. 2009). Domesticated cassava further separated into two groups, sweet and bitter cassava, based on root cyanide content (Elias et al. 2004; Clement et al. 2010). Sweet cassava is generally cultivated throughout the Neotropics, but dominates the western and southern water-heads of the Amazon river basin, while the northern water-heads and central portion are dominated by bitter cassava (Renvoize 1972; Fraser et al. 2012; Mühlen et al. 2019), which is predominantly used for starch extraction and cassava flour production. In both cases, cassava is valued for its starchy storage roots, especially by smallholder farmers. The objective of this study is to provide a comprehensive genetic characterization of Embrapa cassava gene bank to further increase genetic gain in Brazil and explore domestication intermediate events.

Genetic gain from breeding in cassava has made little progress over the last century compared to other crops (Ceballos et al. 2004). Many challenges confronting cassava breeding (limiting breeding efficiency) include its heterozygous genome, long breeding cycles, clonal propagation, and non-recovery of recurrent genome after single trait introgression (Ceballos et al. 2016; Kuon et al. 2019). While cassava is mostly clonally propagated, it is an outcrosser with plants still capable of sexual reproduction, and the conscious or unconscious inclusion of seedlings into clonally propagated stock is continually generating new genotypes within a population, thereby increasing allelic variation (McKey et al. 2010). Modern breeding activities have led to major changes, notably in Africa with *M. glaziovii* introgressions, in key plant architecture traits (Nichols 1947; Lef□vre and Charrier 1993; Wolfe et al. 2019); however, the presence of introgressed segments has not been explored in Brazilian germplasm from past breeding activities. Crop selection is usually extended to adaptive alleles, leading to a reduction in the effective size of a breeding population (Wang, Santiago, and Caballero 2016). Hence, exploring available genetic diversity from Brazilian germplasm collections and wild relatives can provide additional adaptive and reproductive traits to breeding populations. This hypothesis is supported by cultivated cassava progenitors (*M. esculenta spp. flabellifolia*) and wild relatives (*M. glaziovii*) having been shown to have higher nucleotide diversity (Ramu et al. 2017; Wolfe et al. 2019). Embrapa cassava germplasm collections cover the 26 Brazilan states and 5 ecoregions, giving rise to potential population structure through various evolutionary forces, including geographic boundaries (A. Miller and Schaal 2005) and domestication events (Alves-Pereira et al. 2018). *In situ* genetic diversity studies have reported population structure to be different across bitter and sweet Brazilian cassava, owing to high haplotype diversity and environmental heterogeneity of cassava growing regions (Alves-Pereira et al. 2018). However, *ex situ* germplasm collection did not reveal such spatial genetic patterns (de Oliveira et al. 2014; H. Y. G. de Albuquerque et al. 2018). Such agro-ecological information is essential for pre-breeding activities and varietal deployment in target regions (Dwivedi et al. 2016).

One pre-breeding activity that relies critically on understanding population structure is the Genome-wide association study (GWAS); spurious associations may result from a biased distribution of alleles, reflecting the underlying genetic structure and relatedness of individuals in a population. Historically, with Brazil as the center and origin of cassava diversity (Kenneth M. Olsen 2004), germplasm sharing between the Amazonas and the central region of Brazil brought about the dispersal of cassava as the natives migrated across regions in Brazil (H. Y. G. de Albuquerque et al. 2019). With a continuous increase in germplasm exchange globally and as well as within Brazil, along with natural and artificial selection, the renaming of germplasm by farmers became almost impossible to track, challenging germplasm management (Kilian and Graner 2012). Moreover, the extent of Linkage Disequilibrium (LD) defines, on average, the required number of SNP markers and mapping resolution in association studies (Flint-Garcia, Thornsberry, and Buckler 2003). LD decay and population structure has been extensively explored in sexually propagated plants including *Arabidopsis thaliana* and maize (Nordborg et al. 2002; Kim et al. 2007), but only recently in clonally propagated crops including potato, sweet potato and African cassava (Stich et al. 2013; Vos et al. 2017; Wadl et al. 2018; Rabbi et al. 2017; Ramu et al. 2017).

This study provides a genome-wide assessment of population structure, genetic diversity, linkage disequilibrium, introgressions, population splits and gene flow events from Brazil’s Embrapa Cassava Germplasm Bank, center of cassava diversity. Taken together, our results will enable a more effective use of germplasm resources for various breeding-related activities, such as marker development, parent’s selection, and gene mapping.

## Materials and Methods

### Plant material

A total of 3,354 accessions from the Cassava Germplasm Bank (CGB) of Brazilian Agricultural Research Corporation, Embrapa Mandioca e Fruticultura, located in Cruz das Almas, Bahia, Brazil (12 400 19” S, 39 060 22” W, 226 m altitude) were used for this study (**Fig. 1a**). The germplasm collection includes cassava landraces and modern breeding lines from various growing regions and ecoregions of Brazil. A subset (1,580) of these 3,354 accessions was previously described in Albuquerque et al. (H. Y. G. de Albuquerque et al. 2018). The entry years for individuals in studied populations ranges from 1962 to 2018. An additional set of 62 South American accessions from the cassava HapMap efforts were included (Bredeson et al. 2016; Ramu et al. 2017). A subset of 1,389 individuals were phenotypically characterized across multi-year trials for root hydrogen cyanide content (HCN) as described by Ogbonna and colleagues (Ogbonna et al., n.d.).

**Figure 1.**
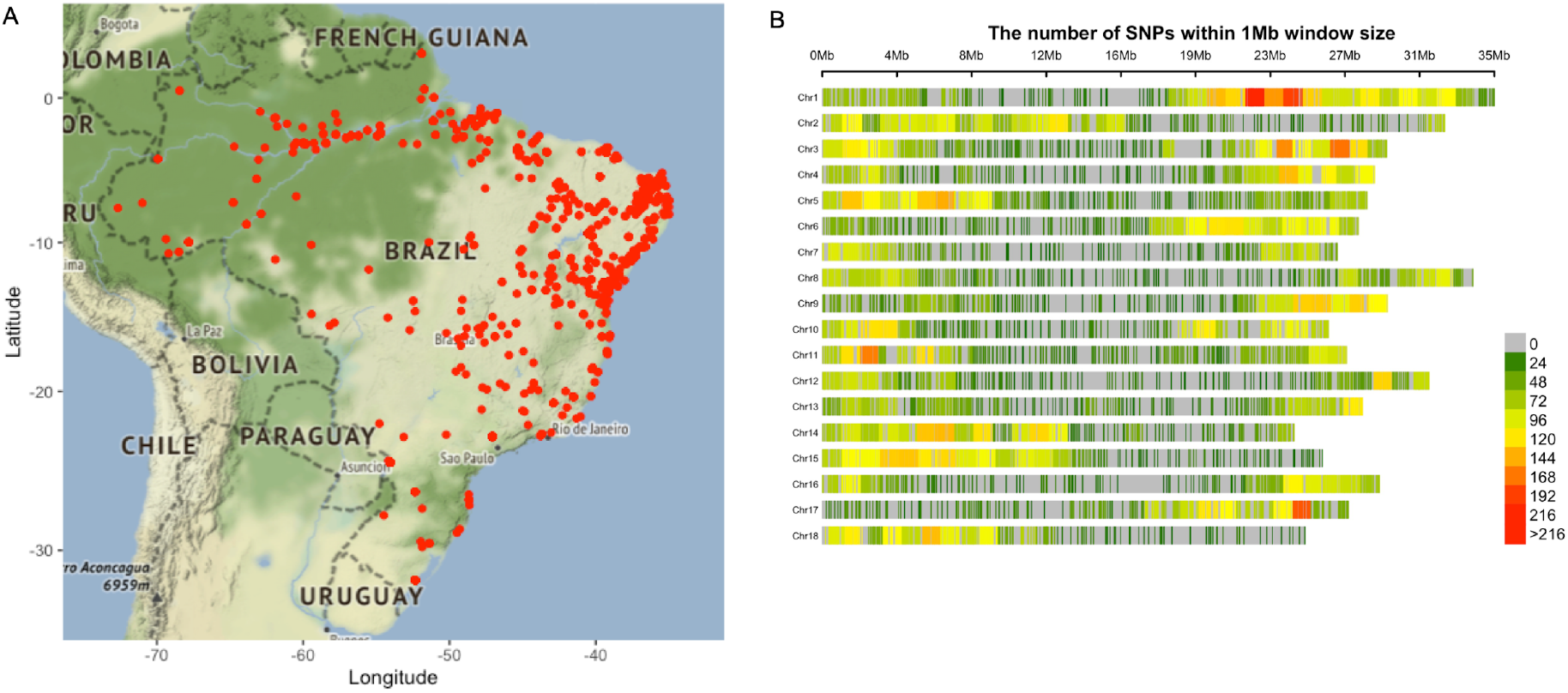
Germplasm geographic sampling and SNP distribution. **(A)** Geographic locations for cassava clones included in the germplasm collection. A total of 1,821 cassava accessions have valid geographic information; each red dot shows the specific site in Brazil where an accession was sampled from. **(B)** Genotyping by sequencing single nucleotide polymorphism (SNP) density in the Brazilian germplasm collection. The legend scale indicates the number of SNPs within 1Mb window size. The plot shows the distribution of SNPs across the genome in our population.

### Datasets

The initial 3,345 germplasm set (annotated GA panel) was subjected to Identity-By-State analysis (see details below) and a unique core set of 1,536 germplasm was constructed (annotated GU panel). To further guide germplasm characterization for pre-breeding activities, 62 known South American cassava landraces and wild relatives from the cassava HapMap (Ramu et al. 2017; Bredeson et al. 2016) were included (annotated GUH and GAH panels). To perform spatial population structure analysis, a total of 1,657 individuals with Global Positioning System (GPS) coordinates were used and annotated GAHg panel. See **Supplementary Table 1** for dataset panels definition.

### DNA extraction

DNA extraction was performed following the protocol described by Albuquerque et al (H. Y. G. de Albuquerque et al. 2018). Briefly, DNA was extracted from young leaves according to the CTAB (cetyltrimethylammonium bromide) protocol as described by Doyle and Doyle (1987), with minor modifications. The quantity of DNA was estimated by comparing the fluorescent yield of the samples with a series of lambda (λ) DNA standards (Invitrogen, Carlsbad, CA) at varying known concentrations. The DNA was diluted in TE buffer (10 mM Tris-HCl and 1 mM EDTA) to a final concentration of 60 ng/μL, and the quality was checked by digestion of 250 ng of genomic DNA from 10 random samples with the restriction enzyme *Eco*RI (New England Biolabs, Boston, MA) at 65°C for 2 hour and thereafter visualised on agarose gel.

### Genotyping

Genotyping was performed by preparing pooled libraries using Genotyping-By-Sequencing (GBS) (Elshire et al. 2011) with the *Ape*KI restriction enzymes (Rabbi et al. 2014) and sequenced on the Illumina HiSeq 2500 platform producing read length of 150 bp. Reads were aligned to the cassava version 6.1 reference genome (Bredeson et al. 2016). Single-nucleotide polymorphism (SNP) calling was performed using TASSEL GBS pipeline V5 (Glaubitz et al. 2014). The SNP calling step included the following parameters: quality: -mnQS 1, kmer length: 64 and the taxa key file. Raw output data was subjected to filtering including: Individual-based missing data filtering of 0.8 maximum per chromosome, mean depth values (over all included individuals) greater than 5, missing data up to 0.2 per locus, and a minor allele frequency of 0.01 per locus. Phasing was performed for each chromosome using beagle4.0 (Browning and Browning 2009), a window of 5,000 markers and an overlap window of 500 markers. Imputation was performed using the genotype likelihood (GL) mode with 10 iteration steps. Imputed Markers were subjected to filtering using an allelic correlation (*AR*^*2*^ > *0.8*) equal or greater than 0.8. Dosage format was generated using the pseq library (http://atgu.mgh.harvard.edu/plinkseq/start-pseq.shtml).

### Population structure and genome-wide relatedness

Population structure analysis was performed using loci satisfying Hardy-Weinberg Equilibrium (--hwe: 0.01) and minor allele frequency (--maf: 0.05) filtering, linkage disequilibrium marker pruning using --indep-pairwise 50 5 0.8 (windows, step, *r*^2^). All filtering was performed using PLINK (v1.9; (Chang et al. 2015) and vcftools (v4.2; Danecek et al. 2011). These GBS-based SNP datasets (i.e., GUH and GAHg) were intersected with the whole-genome sequencing (WGS)-based HapMap dataset (available at ftp://ftp.cassavabase.org/HapMapII/rawData/241_accessions/) using the GATK CombineVariants and SelectVariants functions (McKenna et al. 2010).

Germplasm duplicate identification was performed using a set of 9,686 SNPs filtered on the basis of genome-wide average proportion of alleles shared between individuals using PLINK IBS estimation approach (Purcell et al. 2007). Secondly, a Ward’s minimum variance hierarchical cluster dendrogram was built from the IBS matrix using the APE v.5.3 package (Paradis 2011) in R (R. Core Team 2015) following the calibration principle approach (Noli, Teriaca, and Conti 2013; Rabbi et al. 2015) and a reference set of 11 known duplicated accessions. The phylogeny tree was constructed from Identity-by-State (IBS) pairwise distances using hierarchical clustering hclust function (“ward.D2” method) in phyclust v0.1-28 R package version 3.6.3 (2020-02-29) and visualized in FigTree v1.4.4 (http://tree.bio.ed.ac.uk/software/figtree/).

Based on the IBS-selected unique core set of 1,536 accessions, population stratification analysis was conducted on the GUH panel and 8,242 SNPs using Admixture (Alexander, Novembre, and Lange 2009) with 5 folds cross validations for values of K 1 through 20 to define the optimal number of clusters and examine patterns of relatedness and sub-ancestry among our population. Non-parametric approaches, such as multivariate principal component analysis (PCA) and associated Discriminant Analysis of Principal Components (DAPC) using Bayesian Information Criterion (BIC), were used for validation using Adegenet package v.2.1.2 in R (Jombart, Devillard, and Balloux 2010; Jombart 2008).

To assess family structure and genetic relatedness within our dataset, Identity-by-Descent (IBD) estimation was carried out using PLINK as previously described by (Bredeson et al. 2016) for cassava. Briefly, estimation of IBD between two diploid organisms can be highlighted by three subclasses (IBD0, IBD1, IBD2) with probabilities summarized by Cotterman coefficients (*Z0*, *Z1*, *Z2*, respectively). IBD0, IBD1 and IBD2 indicate the sharing of zero haplotypes (all four haplotypes distinct), one haplotype (two haplotypes unshared) and two haplotypes (all four haplotypes shared) over a defined genomic region for the two individuals.

### Genetic diversity analysis

Observed heterozygosity and average polymorphism information content (PIC) for the identified genetic groups was computed on 27,045 SNPs using R (R. Core Team 2015), according to the methods earlier described by Botstein (Botstein et al. 1980), using the following equation: *PIC* = 1 − *Σ*_*i = 1*_*^n^P_i_^2^* − *Σ*_*i = 1*_ ^*n−1*^ *Σ*_*j* = *i + 1*_ ^*n*^*2P_i_*^*2*^*P*^*2*^_*j*_, where *P_i_* and *P_j_* are the frequencies of and *i*^*th*^ and *j*^*th*^ alleles and *n*is the number of different alleles for any selected SNP. Observed heterozygosity was estimated using radiator software version 1.1.4 R package (Gosselin T. 2019). Minor allele frequency (MAF) were computed using PLINK v2.0 (Purcell et al. 2007), while fixation index (*F_ST_*) was estimated using Weir and Cockerham’s estimator (Weir and Cockerham 1984). Observed heterozygosity was estimated using GENEPOP software version 4.7.5 (Rousset 2008) And effective population size 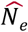, was estimated at an allele frequency of 0.02 using LDNe method (Waples and Do 2008), as described by (Sovic et al. 2019) and implemented in the software NeEstimate version 2 (Do et al. 2014). To limit individual group size incidence on parameter estimates, all of the population genetic parameters were computed on 120 individuals randomly sampled from each identified group (1, 2, 5,–10; **Supplementary Table 2)**. Genetic groups (3, 4) with sample size *n* < 100 were not considered in subsequent downstream analysis.

### Linkage disequilibrium and introgression analyses

Pairwise LD using all markers (27,045 SNPs) with a minor allele frequency (MAF) 5% on each chromosome was computed using PLINK. In addition, genome-wide LD decay was computed for each identified genetic group. To limit individual group-size incidence on LD estimates, 120 individuals were randomly sampled from each group, as earlier mentioned, for population genetics parameter estimations and LD decay curves were fitted using non-linear regression following previously described approaches (Van Inghelandt et al. 2011; Bauchet et al. 2017).

To identify potential interspecific hybridization between cultivated (*M. esculenta* ssp. *esculenta*) and wild material (*M. esculenta* ssp. *flabellifolia, M. glaziovii*), introgression detection analysis fo*r M. esculenta* ssp. *flabellifolia* and *M. glaziovii* was carried out using the method earlier described by Bredeson and colleagues (Bredeson et al. 2016; Wolfe et al. 2019). Here, we used the whole-genome sequencing HapMap dataset (Ramu et al. 2017) and contrasted groups (cultivated and progenitors) of 7 representative accessions each based on the phylogenetic analysis from Ramu et al. (Ramu et al. 2017), see **Supplementary Table 3**.

### Spatial population structure, and population splits, and genome scan for selection

To further investigate the relationship between genetic clusters and available geospatial coordinates, we computed ancestry estimates for the GAHg panel (1,657 individuals) and a genome-wide scan using TESS3 (Caye et al. 2016). Briefly, TESS3 is a model-free approach which performs spatial ancestry estimation using least-squares optimization and on geographically constrained non-negative matrix factorization. Ancestral genotype frequency matrix G and ancestral coefficient matrix Q are used to derive the allele frequencies in the K ancestral populations. Locus-specific *F_ST_* statistics are computed based on the estimated ancestral allele frequencies. The *F_ST_* statistics are transformed into squared *z*-scores and p-values are computed using a Chi-squared distribution with K − 1 degrees of freedom (Weir 1997), where K is the number of ancestral populations (Caye et al. 2016). A set of 1,039 common accessions between GAHg and GUH panels were used (**Supplementary Table 1**) to compare results between non-spatial (DAPC) and explicitly spatial (TESS3) population structure analysis.

To infer intermediate evolutionary stages of domestication, including population splits and migration events, we used 27,045 SNPs and 419 Brazilian germplasm (**Supplementary Table 4**) collected before the year 2000 (contemporary breeding lines excluded). We constructed a consensus population graph tree using SNP window size of 50, five migration events with sample correction turned off, and 1000 bootstrap replications. The TreeMix maximum likelihood method (Pickrell and Pritchard 2012), as implemented in the BITE R package version 1.2.0008 (Milanesi et al., n.d.), was used to highlight graph robustness. The approach is based on population genomics using genome-wide allele frequency and Gaussian approximation to genetic drift.

## Results

### Genotyping-by-sequencing and marker distribution

From the SNP calling, a total of 343,707 initial variant loci were identified. Of these, 30,279 were retained for phasing and imputation after filtering for quality, missing data, and minor allele frequency. After imputation, 27,045 SNPs with an allelic correlation of *AR*^*2*^≥ 0.8 were kept for downstream analysis. Genotyping density is, on average, one SNP every 19 kilobase pairs (kbp), with a maximum distance of 2 megabase pairs (Mbp). Marker density per chromosome, ranges between one SNP every 14 kbp (chromosome 1) and one SNP every 27 kbp (chromosome 7), with an average of 1,503 markers per chromosome (**Fig. 1b**). For IBS analysis on the GA panel, 9,686 SNPs remained after Hardy-Weinberg filtering, while 8,242 SNPs remained after additional minor allele frequency filtering on the IBS-selected unique set (GU).

### Duplicate accession identification and population structure

Because biased representation of duplicated individuals in a dataset will lead to downstream biases in the estimated population genetics parameters, we conducted an identity-by-state (IBS)-based duplicate analysis. This analysis yielded 1,536 unique clusters [with Ward’s distance threshold of 0.004] (**Fig. 2a**, **Supplementary Table 5, Supplementary Figure 1**) and 1,818 duplicates. Based on available metadata, the duplicates are distributed across the five ecoregions, with 85% coming from the Northeastern region, covering Cerrado, Caatinga and Atlantic Forest ecoregions, while 13% and 2% comes from the Amazonas and Pampa ecoregions, respectively.

**Figure 2.**
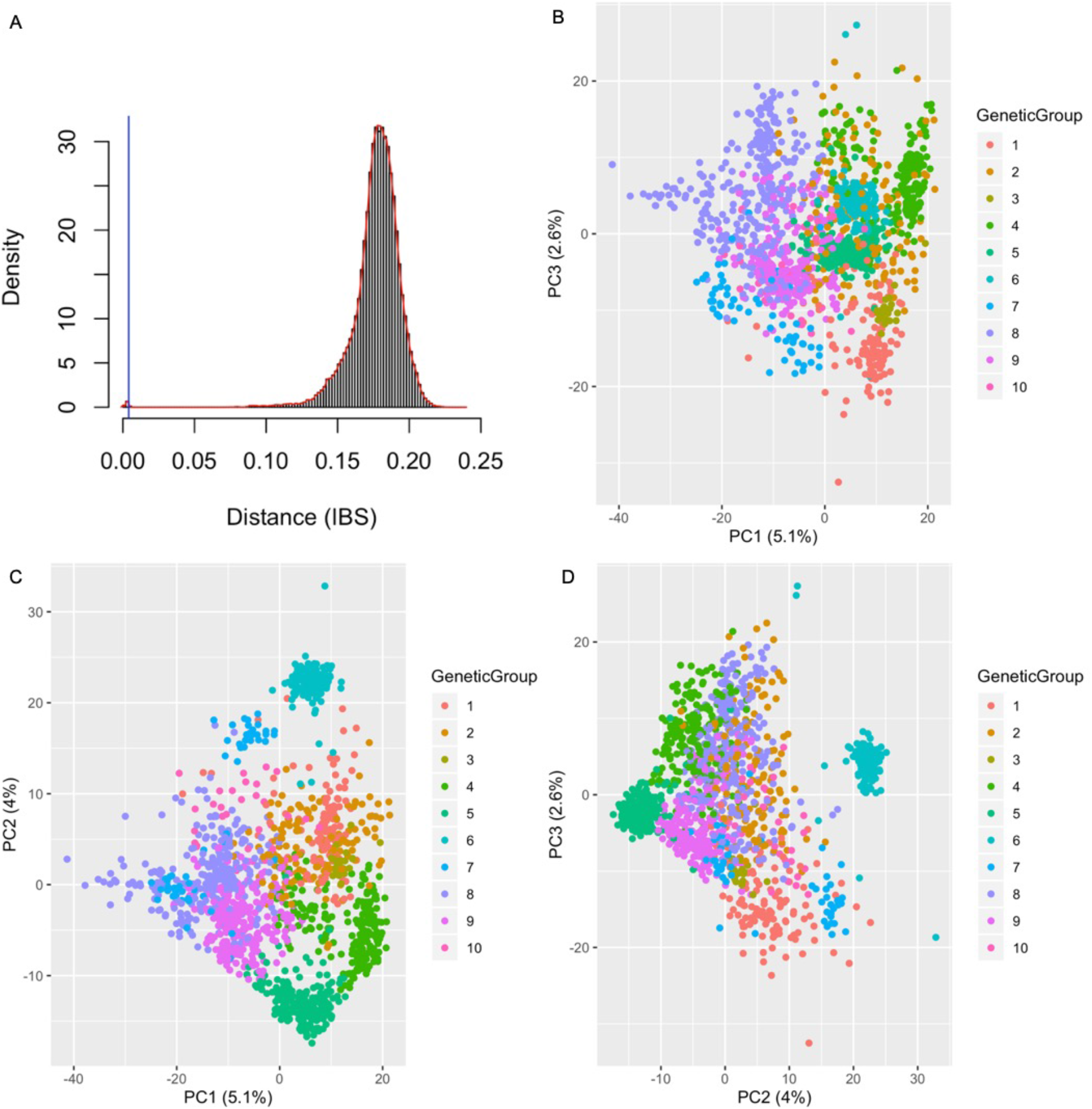
Identity by State analysis and Population structure. **(A)** Density plot of IBS values: the left tail shows values close to 0. In IBS scaling, 0 equals identical accessions. The average IBS value for these duplicate accessions was **0.003135969** after excluding the artefacts. Based on a calibration set we considered the conservative threshold of 0.004, yielding 1,818 entries involved in duplication. The blue vertical line denoting the distance threshold below which two samples can be considered identica. **(B-D)** Principal Component Analysis of GU (IBS-selected unique set) dataset. Population structure of Brazilian germplasm reveals the first three axes of the principal component analysis (PCA) explain about 11.1% of the variations in the population of 1,536 individuals (GU panel), which includes 8,242 SNPs after filtering for Hardy-Weinberg equilibrium and minor allele frequency of 0.01 and 0.05 respectively. (B) Plot of PC1 and PC3 explaining 7.7% of the variations. **(C)** Plot of PC1 and PC2 explaining 9.1% of the variations. **(D)** Plot of PC2 and PC3 explaining 6.6% of the variations. The color codes represent the genetic groups identified using the DAPC of GUH panel (IBS-selected unique set plus 62 HapMap accessions from Brazilian germplasm).

To further dissect the genetic structure of our dataset, PCA, DAPC, and Admixture analyses were conducted. Non-parametric methods such as PCA and DAPC can provide accurate population structure estimates in complex scenarios (ie: continuous admixture, complex substructure) and lead to improved population inference. Admixture models, however, are more interpretable when using spatial information (François and Durand 2010). When combined, these approaches provide a platform for understanding the spatial distribution of adaptive and nonadaptive genetic variation in different organisms (Manel et al. 2010). PCA revealed patterns of clustering in our population, with the first three PCs accounting for 11.7% of the genetic variation (**Fig. 2b–d**). To ensure optimal statistical power for evaluating genetic structure, and obtaining the most accurate number of clusters using DAPC, we kept 500 PCs that explained about 90% of the genetic variance in our dataset (**Supplementary Figure 2**), following observations made by Jombart and colleagues (Jombart, Devillard, and Balloux 2010). Application of the Bayesian Information Criterion (BIC) identified an optimal number of ten genetic clusters in the GUH panel (**Fig. 3b**, **Supplementary Figure 3a–b**), with group size ranging from 25 to 265 individuals (**Table 1**). The probability of individual group assignment was 99%, showing stability, and indicating that 10 clusters adequately summarizes our dataset. The DAPC group identification was further validated using Admixture, a parametric approach with five fold cross-validation, also identifying ten subpopulations (**Figure 3a**, **Supplementary Figure 3c–e**). Further analyses and estimation of population genetics parameters were performed on the genetic groups identified in the GUH panel.

**Figure 3.**
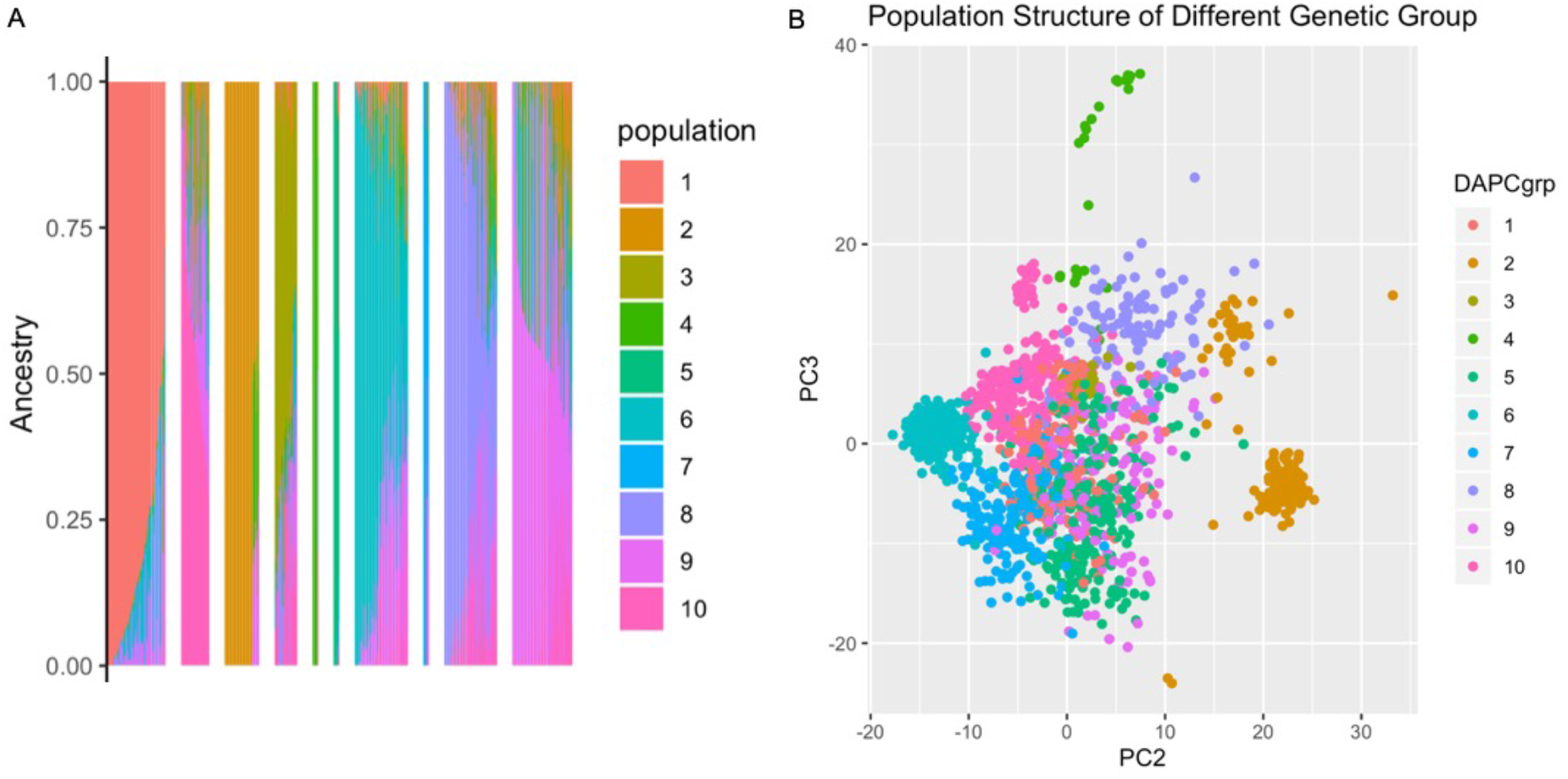
Admixture analysis and Discriminant Analysis of Principal component for GUH dataset with 1,536 unique sets and 62 HapMap individuals. **(A)** ADMIXTURE analysis (K = 10) with the probability of assignment ordered sequentially. Proportion of ancestry was plotted on the Y-axis against the individual samples on the X-axis. **(B)** GUH dataset PC2 and PC3 view of population structure with individuals colored by their DAPC genetic groups.

**Table 1:**
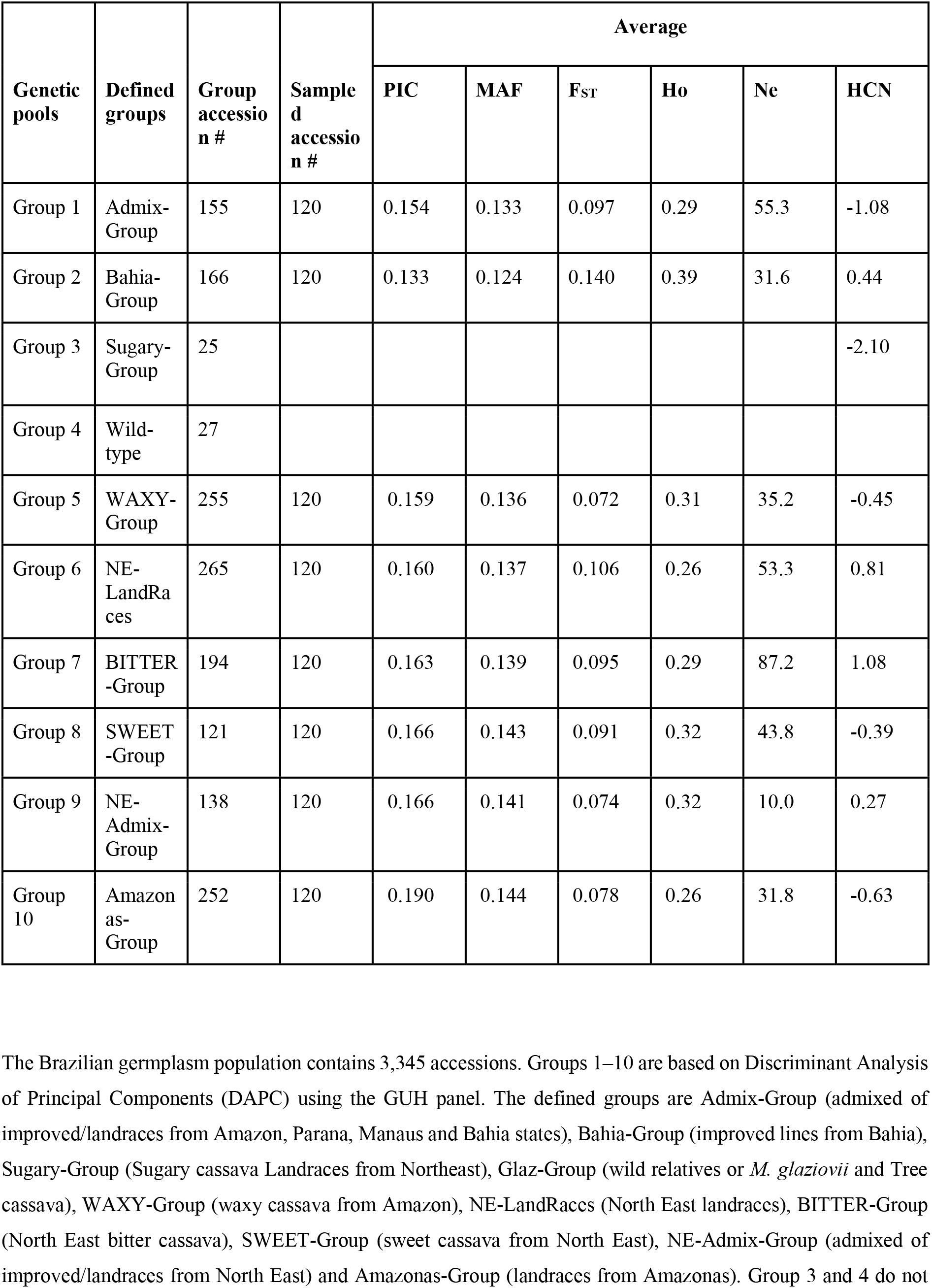
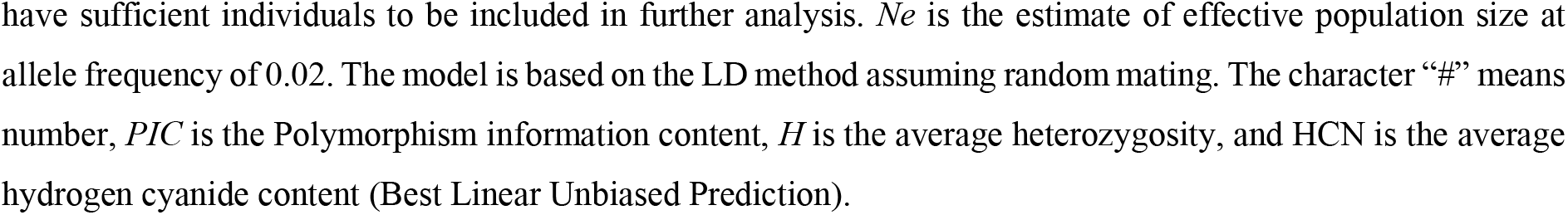
Genetic diversity of Brazilian germplasm.

| Genetic pools | Defined groups | Group accession # | Sample d accession # | Average |  |  |  |  |  |
| --- | --- | --- | --- | --- | --- | --- | --- | --- | --- |
|  |  |  |  | PIC | MAF | F<sub>ST</sub> | Ho | Ne | HCN |
| Group 1 | Admix-Group | 155 | 120 | 0.154 | 0.133 | 0.097 | 0.29 | 55.3 | -1.08 |
| Group 2 | Bahia-Group | 166 | 120 | 0.133 | 0.124 | 0.140 | 0.39 | 31.6 | 0.44 |
| Group 3 | Sugary-Group | 25 |  |  |  |  |  |  | -2.10 |
| Group 4 | Wild-type | 27 |  |  |  |  |  |  |  |
| Group 5 | WAXY-Group | 255 | 120 | 0.159 | 0.136 | 0.072 | 0.31 | 35.2 | -0.45 |
| Group 6 | NE-LandRaces | 265 | 120 | 0.160 | 0.137 | 0.106 | 0.26 | 53.3 | 0.81 |
| Group 7 | BITTER-Group | 194 | 120 | 0.163 | 0.139 | 0.095 | 0.29 | 87.2 | 1.08 |
| Group 8 | SWEET-Group | 121 | 120 | 0.166 | 0.143 | 0.091 | 0.32 | 43.8 | -0.39 |
| Group 9 | NE-Admix-Group | 138 | 120 | 0.166 | 0.141 | 0.074 | 0.32 | 10.0 | 0.27 |
| Group 10 | Amazonas-Group | 252 | 120 | 0.190 | 0.144 | 0.078 | 0.26 | 31.8 | -0.63 |

The Brazilian germplasm population contains 3,345 accessions. Groups 1–10 are based on Discriminant Analysis of Principal Components (DAPC) using the GUH panel. The defined groups are Admix-Group (admixed of improved/landraces from Amazon, Parana, Manaus and Bahia states), Bahia-Group (improved lines from Bahia), Sugary-Group (Sugary cassava Landraces from Northeast), Glaz-Group (wild relatives or *M. glaziovii* and Tree cassava), WAXY-Group (waxy cassava from Amazon), NE-LandRaces (North East landraces), BITTER-Group (North East bitter cassava), SWEET-Group (sweet cassava from North East), NE-Admix-Group (admixed of improved/landraces from North East) and Amazonas-Group (landraces from Amazonas). Group 3 and 4 do not
have sufficient individuals to be included in further analysis. $N_e$ is the estimate of effective population size at allele frequency of 0.02. The model is based on the LD method assuming random mating. The character “#” means number, $PIC$ is the Polymorphism information content, $H$ is the average heterozygosity, and HCN is the average hydrogen cyanide content (Best Linear Unbiased Prediction).

To further understand the observed population structure, all individuals were mapped to their available metadata, providing insights into the observed clustering pattern (see **Supplementary Table 6**). Clustering approaches (Admixture, PCA, DAPC) and metadata on the GUH dataset supported their group identification (See **Table 1**). **Group 1**, with 155 accessions, is an admixed population of landraces and improved breeding lines from Amazonas, Bahia, Parana and some exchange materials from Manaus (annotated **Admix-Group**). **Group 2**, with 166 admixed accessions, is made up of family-structured improved lines from Bahia (annotated **Bahia-Group**); while **group 3**, with 25 accessions, contains landraces of sugary cassava from Northeastern Brazil (annotated **Sugary-Group**); **group 4**, with 27 accessions, is highly correlated with *M. glaziovii* and made up of materials only from HapMap accessions from Brazil (annotated as **Glaz-Group**); while **group 5**, with 255 accessions, is an admixed population and made up of landraces and improved lines including waxy clones from the Embrapa breeding program (annotated as **WAXY-Group**). **Groups 6 and 7**, with 265 and 194 accessions, respectively, are cassava accessions from the North Eastern region of Brazil and are made up of landraces and bitter cassava respectively (annotated as **NE-LandRaces** and **Bitter-Group**). **Group 8**, with 121 accessions, is made up of sweet cassava from North Eastern Brazil (annotated as **SWEET-Group**). **Group 9**, with 138 accessions, is an admixed population of landraces and improved breeding lines from North Eastern Brazil (annotated as **NE-Admix-Group**); while **group 10,** annotated as **Amazonas-Group** with 252 accessions, is made up of landraces from the Amazonas and clustered with the *M. esculenta* ssp. *flabellifolia* from the HapMapII accessions (**Fig. 3a–b)**. Average root cyanide content per group is shown in **Supplementary Figure 3f,** with Sugary-Group and Bitter-Group having the lowest and highest of cyanide respectively (Ogbonna et al., n.d.).

### Genetic diversity parameters

The average probability of individual assignment into the different DAPC genetic groups was 99% with a maximum of 265 accessions (21% of GUH panel) assigned to group 6 and a minimum of 25 accessions (2%) assigned to group 3 (**Table 1**). A limited sample size of a population can affect population genetic parameter estimates (Bashalkhanov, Pandey, and Rajora 2009) hence, groups 3 and 4 (25 and 27 accessions) were not included in further population genetics estimation analysis. Bahia-Group, the family-structured-group was also excluded from the reported genetic diversity parameters.

Minor allele frequency (MAF) across groups showed an average of 0.14, Amazonas-Group and Admix-Group having the highest (0.144) and lowest (0.133) MAF, respectively (**Fig. 4a**, **Table 1)**. Group pairwise *F_st_*(**Fig. 4b**) ranged from 0.037 (Admix-Group and WAXY-Group) to 0.122 (Admix-Group and Bitter-Group), with NE-LandRaces (0.105) having the highest differentiation when compared to the rest of the populations. The overall *F_st_* was 0.079.

**Figure. 4.**
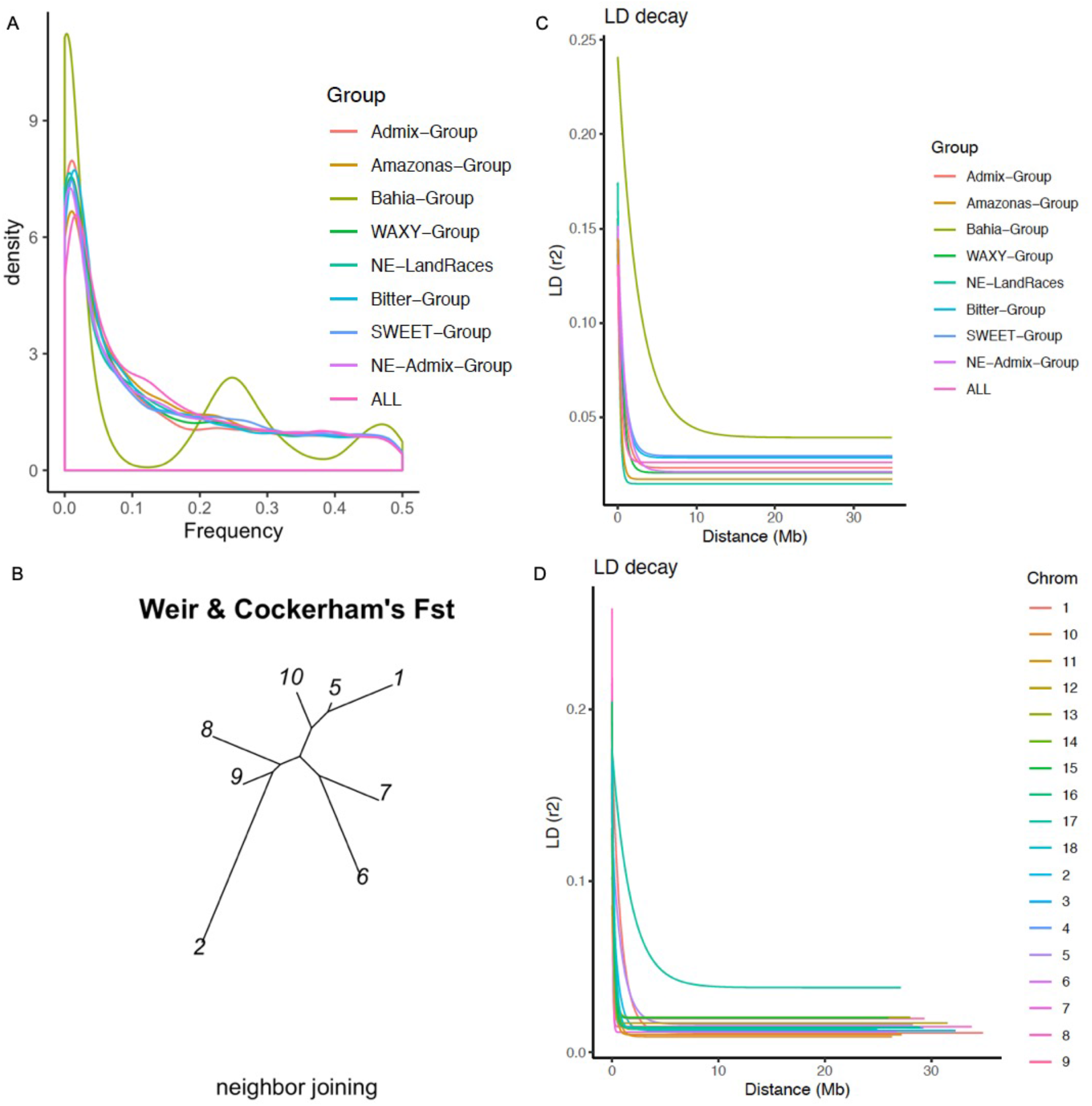
Genetic parameters and linkage disequilibrium on GUH panel. **(A)** Minor allele frequency overview across identified genetic groups and combined groups denoted as “ALL”. **(B)** Neighbor joining tree showing pairwise Fst of the identified groups in Brazilian population. The groups are 1, 2, and 5 through10 (respectively: Admix-Group, Bahia-Group, WAXY-Group, NE-LandRaces, Bitter-Group, SWEET-Group, NE-Admix-Group, and Amazonas-Group). **(C)** Linkage disequilibrium decay based on Hill and Weir model (*r²*) for each identified genetic group and all groups combined accessions, plotted along physical distances across the genome and along the genetic distances in megabase (Mb). Across the whole population is denoted as “ALL” in the legend. **(D)** Trend line of the non-linear regression of the linkage disequilibrium measure r^2^ versus physical distance (Mb) between single nucleotide polymorphism (SNP) marker pairs across chromosomes in the genome of the Brazilian germplasm. The LD is computed chromosome-wide. The average LD r^2^ is 0.031 and drops to background level (r^2^< 0.1) at around 300 kb across the genome.

The average PIC across groups was 0.23 (max = 0.375; min = 0.142; **Table 1**). Approximately, 25% of the SNPs have PIC values greater than 0.3, while 42% had PIC values greater than 0.2 for the population of 3,345 accessions. PIC values for identified groups ranged from 0.190 to 0.154, with an average of 0.165. Admix-Group (0.154) showed the least genetic diversity and Amazonas-Group the highest PIC value (0.190) compared to the rest of the groups and also higher than the PIC value of the combined genetic pool. The observed heterozygosity (*H_o_*) across groups ranges from 0.26 (Amazonas-Group) to 0.32 (SWEET-Group and NE-Admix-Group) with an average of 0.29 (**Table 1**). While estimates of effective population size ranges from 10.0 (NE-Admix-Group) to 87.2 (Bitter-Group), as seen in **Table 1**.

### Linkage Disequilibrium and Introgression Analysis

Average genome-wide LD was 0.021 (*r^2^*) and drops to background level (*r^2^* < 0.1) at around 107 kb (**Fig. 3c; Supplementary Figure 4, Supplementary Figure 5, Supplementary Figure 6**). All identified groups and chromosomes exhibited average LD (*r^2^*) less than 0.1 at genome-wide scale (**Supplementary Figure 5a, Supplementary Table 7).** Higher chromosomal LD extent was observed on chromosome 17 with *r^2^* value of 0.06 (**Supplementary Figure 4, Supplementary Figure 5a**), and slowest decay (**Fig. 4d**). Average LD across the genome, excluding chromosome 17, is 0.019 (*r^2^*). Bahia-Group, a family-structured-group, excluded from the LD analysis reporting, showed a much higher physical distance of 2.98 and 4.15 Mb for *r^2^* > 0.1 and *r^2^* > 0.083, respectively (see **Supplementary Table 7; Supplementary Figure 5b; Fig. 3c**). An additional description of Bahia-Group is in **Supplementary Note 1** and **Supplementary Figure 7**.

The LD interval size estimates for regions with *r^2^* > 0.1 exhibited variations between groups, ranging from 0.12 Mb, in Amazonas-Group, to 0.4 Mb, in NE-Admix-Group, with 0.13 Mb on average. While the interval size estimates for average LD across groups ranged from 0.86 Mb, in NE-LandRaces, to 2.2 Mb, in NE-Admix-Group, and 1.43 Mb on average (**Supplementary Figure 5b**). At chromosome level across the groups, LD decay ranged from 0.05 Mb in Amazonas-Group to 3.87 Mb in SWEET-Group (**Supplementary Figure 6b, Supplementary Table 7**). The estimated number of SNP markers required for association studies, assuming an LD decay per chromosome with *r^2^* threshold 0.1, ranged from 1,296 to 4,713 SNP markers, (average LD, 236 to 603 SNP markers) and showed broad variation across chromosomes and genetic groups (**Supplementary Table 7**).

Introgression analysis using cassava HapMap reference lines for cultivated *M. esculenta* ssp. *esculenta* vs both wild *M. esculenta* ssp. *flabellifolia* and *M. glaziovii* samples, identified 294 (**Supplementary Table 8**) and 1,795,898 (**Supplementary Table 9**) bi-allelic, ancestry-informative SNP markers, respectively, distributed across chromosomes that represent fixed, or nearly fixed, differences between cultivated and wild cassavas (**Supplementary Figure 8a–b)**. Of the *M. glaziovii*-specific SNPs, 3,238 are represented in the GBS marker set (**Supplementary Figure 8c; Supplementary Table 10)**, which were used to infer ancestral segments in the identified genetic groups. Introgression of *M. glaziovii* in the Brazilian Germplasm collections was found on two chromosomes. The largest introgressed segment (16.8 Mb size, in interval 458800–17307782 bp) was found in chromosome 17, and mainly in Bitter-Group (**Fig. 5**). Bahia-Group and SWEET-Group showed a smaller segment (1.1 Mb size, in interval 5755106–6884548 bp) in chromosome 5 (**Supplementary Figure 9**).

**Figure 5.**
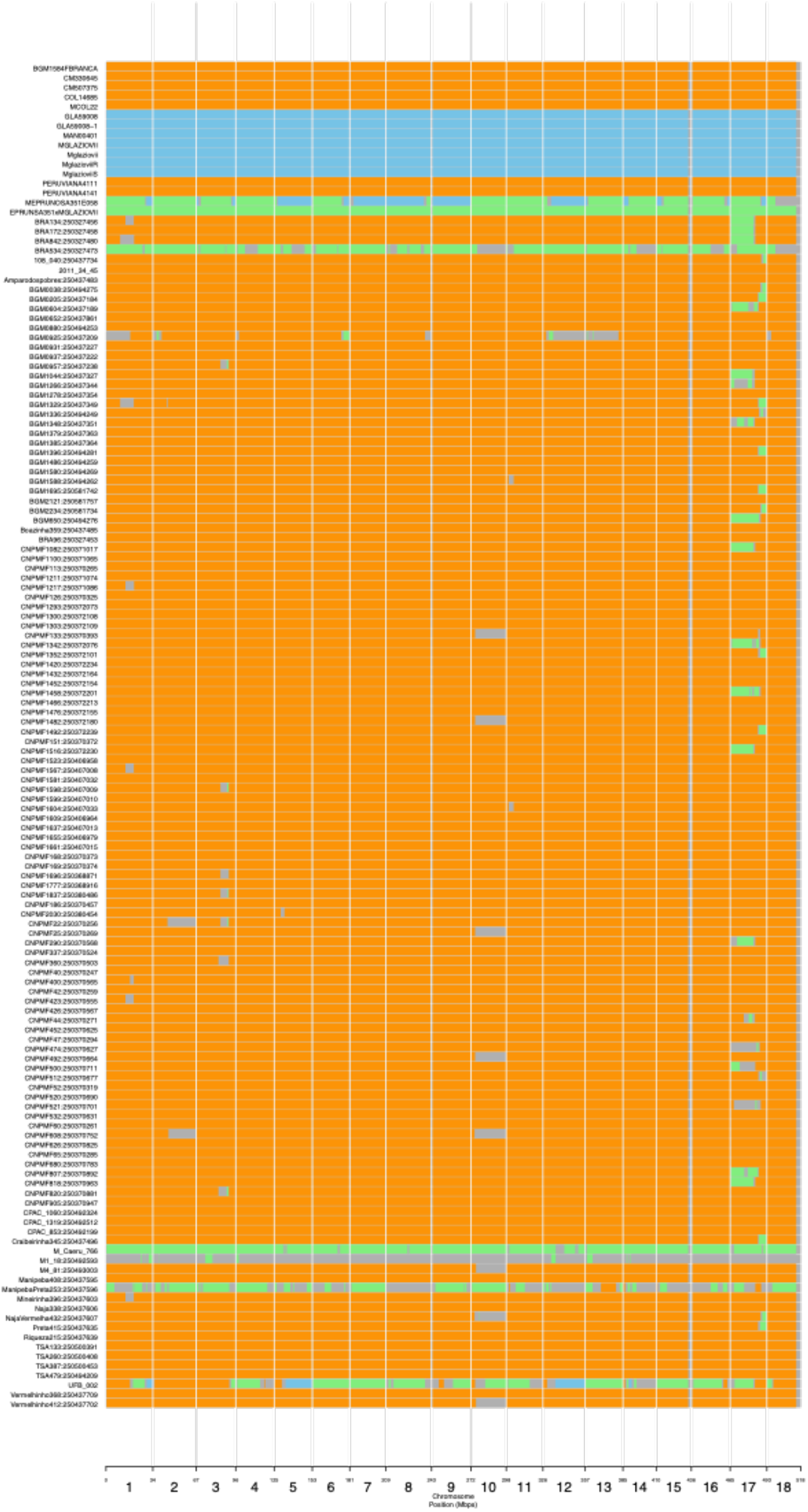
Inferred segmental ancestry of genetic Bitter-Group from Brazilian germplasm collection using Genotyping-By-Sequencing. Bitter-Group is made up of bitter cassava landraces from the Northeastern region of Brazil. Orange color shows both diploid haplotypes are *M.esculenta*; blue is both haplotypes *M.glaziovii*; green, heterozygote haplotypes from *M. esculenta and M. glaziovii*; grey color represent unassigned segments.

### Spatial population structure and genome scan for selection

The number of ancestral populations, *K*, was chosen after evaluating the cross-entropy criterion curve (Frichot et al. 2014), which exhibited a monotonic decrease and plateaued at *K* = 9 (**Supplementary Figure 10a)**. Spatial analysis detected five major ancestral populations and nine extent populations (**Supplementary Figure 10a)**. The nine clusters detected in this spatial analysis as against the ten clusters detected using DAPC and Admixture, was as a result of missing individuals in Glaz-Group, given that they had no GPS coordinates. We show the interpolation of ancestral populations of K = 4 through K = 9 (**Supplementary Figure 10b)**. Between *K*= 4 and *K*= 5, an elbow in the curve was observed (*K*= 5, green line). For *K* greater than 9, cross-validation scores decayed until *K*= 20. Corresponding ancestry coefficients estimate (shown in a biplot; **Supplementary Figure 10b)** interpolation on a geographic map of Brazil were produced for *K*= 4 to 9 ancestral groups (**Supplementary Figure 10c)**. Evaluating the interpolated ancestry at *K*= 4, individuals from Caatinga and Cerrado ecoregions grouped together. At *K* = 5 through 7, five stable groups were resolved; the individuals from Caatinga and Cerrado differentiated into two ecoregions and individuals in the Pampa ecoregion grouped together with Cerrado (**Supplementary Figure 10c, Fig. 6a**). At K = 8 and *K***= 9**, there were additional differentiations (two zones) within the Cerrado and the Atlantic Forest ecoregions. (**Supplementary Figure 10c, Fig. 6a**).

**Figure 6.**
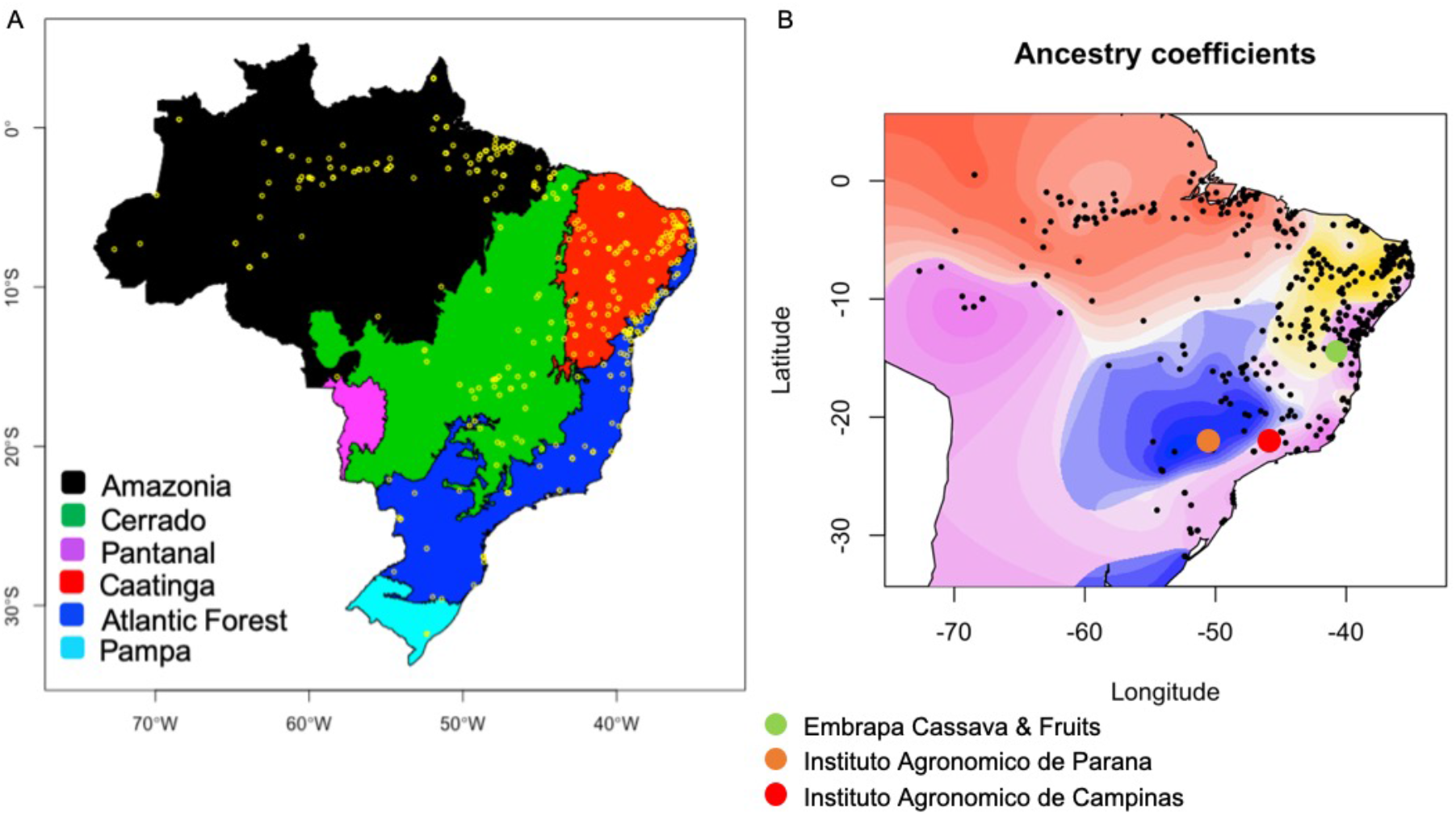
Brazilian ecoregions and Spatial analysis of GAHg panel (1,657 accessions) using tess3r. (A) Map of Brazilian ecoregions highlighting the different biomes, including yellow dots representing the germplasm collection-point coordinates. (B) Geographic maps of ancestry coefficients for K = 5 ancestral populations. The black dots represent the germplasm collection points in Brazil. The fifth cluster is not strongly localized, hence hidden (see **Supplementary Figure 13** for ancestral population coefficient localization). Colored dots (red, orange, and green) represent the geographical locations of the main Brazilian breeding programs active in the past 40 years.

Interestingly, the interpolated ancestry resulting from the five stably-formed groups (from *K* = 5 through 7) present strong geospatial overlap with the six Brazilian ecoregions (*Biomas e sistema costeiro-marinho do Brasil: compatível com a escala 1:250 000* 2019); this excludes the Pantanal ecoregion, where no samples were collected for this study (**Supplementary Table 11**). In **Fig. 6b**, the region in yellow/shades is associated with the semi-arid region of the Brazilian Northeast with shallow and stony soil, sparse vegetation, and low levels of annual rainfall, while the region in red/shades represents the coastal region with high annual rainfall, deeper soils, with higher and dense vegetation. The fifth cluster in **Fig. 6b** is not strongly localized, hence was hidden, thereby displaying only four clusters/colors of ancestral population coefficients. Using 702 landraces (excluding breeding lines), collected before the year 2000, distributed across the five ecoregions of Brazil, the genetic groups show strong overlap with the ecoregions (Chi-squared statistics: X-squared = 593.65, df = 12, p-value < 2.2e–16) (**Supplementary Figure 11, Supplementary Table 12**). A genome-wide scan for selection, based on population differentiation in the nine ancestral populations, detected 24 loci above the Bonferroni-adjusted threshold (**Supplementary Figure 12, Supplementary Table 13)**. The average percentage of accessions grouped in the same groups between GUH panel DAPC and spatial analysis GAHg panel was 85% and ranged between 96% to 53% with only two groups having less than 86% (Bitter-Group: 77% and SWEET-Group: 53%), given that GAHg panel contains 1,036 individuals in common with GUH panel (**Supplementary Table 14**).

TreeMix gene flow analysis showed three population splits with node robustness values of 1000, 636, and 961, respectively, out of 1000 bootstrap replications. Three migration events were thus identified with 0.999 variance explained, indicating the goodness-of-fit of the model with gene flow from Amazonia to Caatinga, Amazonia to Cerrado, and Pampa to Atlantic Forest ecoregions. The Atlantic forest ecoregion had the least genetic drift, followed by Amazonia and Caatinga, and then Pampa and Cerrado ecoregions, respectively (**Fig. 7**).

**Figure 7.**
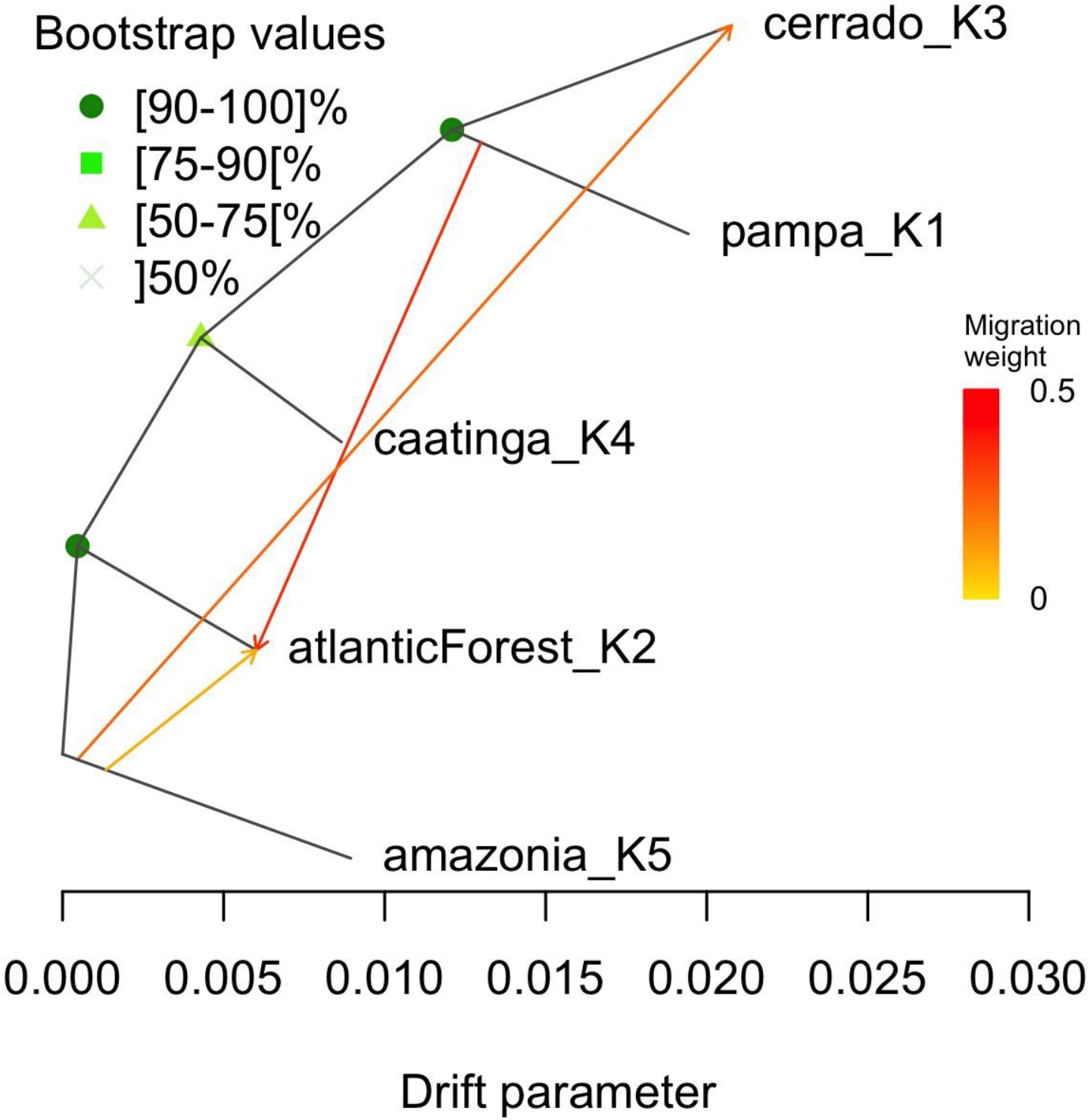
Gene flow analysis using TreeMix (v1.12) for 419 Brazilian germplasm collected before the year 2000. Plotted is the structure of the graph inferred with gene flow events of 3 migration using Maximum likelihood. Individuals in the 5 populations from 5 different Brazilian Ecoregions. Migration arrows are colored according to their weight. Horizontal branch lengths are proportional to the amount of estimated genetic drift. Nodes robustness was estimated with 1000 bootstrap replicates. The analysis was carried out under the implementation of BITE R package version 1.2.0008.

## Discussion

Using 27,045 SNP markers obtained from the Genotyping-by-sequencing (GBS) method, we identified and characterized genetic groups of the Embrapa Brazilian cassava germplasm collection, and inferred the parameters describing genetic diversity, population structure, and linkage disequilibrium.

### IBS Duplicate Identification

Accession mislabelling has been reported to reduce the genetic gain of cassava by up to 40% in Africa (Yabe, Iwata, and Jannink 2018). In order to address germplasm duplication in the Brazilian collection, we inferred genetic identity between accessions using an identity-by-state (IBS) analysis. Pervasive identity between accessions was indeed observed among the Embrapa collection, identifying 1,818 (54%) duplicates out of the 3,354 initial accessions examined in this study. The main outcome of the IBS analysis is that (1) Unique set of 1,536 accessions were identified (GU panel) using Ward’s distance threshold of 0.004 (**Fig. 2a**, **Supplementary Table 1**). (2) This would reduce biases from over representation and genetic contribution of duplicated individuals to genetic variance in a population. Our motivation in this study for IBS analysis was to inform efficient germplasm and resource management at the Brazilian cassava program and balance individual genetic contribution to population structure definition. About 85% of the duplicated individuals were found within the Northeastern region, this region hosts the three cassava breeding programs in Brazil, highlighting the impact of germplasm exchange and adoption of improved varieties by farmers, which are often renamed.

### Population Genetic and Geospatial Structure

To better characterize individuals in our population, we mined common loci between our GBS dataset and 62 WGS HapMap individuals from Brazil, creating the 1,536 accession GUH dataset. The observed DAPC structure patterns of 10 groups, corresponding to the stepping stone model as described by Jombart et al. (2010), which was fewer than previously reported by (de Oliveira et al. 2014; H. Y. G. de Albuquerque et al. 2018). From these previous studies in Brazil, some of the genetic groups reportedly had less than ten accessions in them, indicating that they share similar genomic regions. However, the germplasm used in this study was collected across the 26 states (including Federal district) and over five ecoregions of Brazil, indicating breeding activities, geographical boundaries, and restrictions to have influenced the observed population structure (**Fig. 1a**, **Fig. 6a**).

The distribution of germplasm and genetic diversity across Brazil is heterogeneous with more diverse groups found within the regions where breeding activities have taken place in the last 40 years, while the Amazonas region had less group distributions compared to the rest of Brazil (**Fig. 6b**, **Supplementary Figure 3b**). This observation was further supported by the ancestral coefficient estimated via Admixture, where germplasm within the Amazonas had less diverse ancestral coefficient than germplasm within the Central-West and Southeastern regions (Cerrado and Atlantic Forest ecoregions) of Brazil (**Supplementary Figure 3d–e**). This indicates that a more diverse population is available in Cerrado, Atlantic Forest and Caatinga than in the Amazonas.

The estimated admixture coefficients provided clear evidence for the clustering of homogeneous genetic groups in Brazil ecotypes/ecoregions. The interpolation and projection of ancestry coefficient on Brazilian map suggested a clinal variation along different biotypes/ecoregions across Brazil with a distinct ecotone between ecoregions. The differentiation at *K* = *5*, indicates a significant past event such as domestication, environmental selection or modern breeding activities leading to the differentiation of individuals initially grouped together at *K* = *4* between the Northeast and Central-West regions (Caatinga and Cerado ecoregions) (**Supplementary Figure 10a**). This hypothesis may be supported by **Fig. 6b**, highlighting regions of breeding activities in the past 40 years outside of where the divergence took place. However, future study should consider performing coalescence analysis among individuals from different ecoregions to confirm if the observed divergence is associated with recent breeding activities or a past event associated with cassava domestication.

The cross-validation criterion value at *K* = *4*, indicates four major genetic groups and three major ancestral populations in the GAHg dataset panel. Recently, Muhlen et al. (2019), hypothesized that there are two groups each of sweet and bitter cassava using a country-wide sample of 494 cassava landraces covering 11 geographic regions of Brazil. In our study, the smaller breakups observed from *K* = *6* through 9 indicate additional breeding activities mostly within the Northeast, Central-West, Southeast and Southern regions (covering Cerrado, Caatinga, Atlantic Forest and Pampa ecoregions), highlighting regions of active cassava breeding in Brazil. Above *K* = *9*, indicated that subtle substructure resulting from complex historical isolation-by-distance processes could also be detected (**Supplementary Figure 10a)**. The nine genetic groups identified were as a result of having only 1,039 individuals from the GUH panel with Global Positioning System (GPS) coordinates. The individuals clustered in 9 groups and matches the groupings in DAPC GUH panel, except for Glaz-Group (group 4) where individuals are wild relatives from Whole-genome sequencing HapMap (Ramu et al. 2017) accessions. The average percentage of matching group accessions was 85% across groups and indicates similarity in group assignment between the two independent structure analysis (**Supplementary Table 14)**. From our spatial analysis, especially at K = 5, we speculate that the significant divergence observed between the Caatinga (Northeast region) and Cerrado - Pampa (Central-West and South regions) ecoregions, may represent the second phase of cassava domestication from initial “sweet” domesticated cassava to “sweet and bitter” cassava (**Fig. 6b**). This hypothesis was earlier suggested by (Arroyo-Kalin 2010) and further supported by (Mühlen et al. 2019) and colleagues (Mühlen et al. 2019). (Mühlen et al. 2019) reported that sweet cassava landraces were dispersed southward through the Cerrado where they formed a genetically distinct group that is clearly different from sweet cassava landraces in Amazonia and coastal Brazil. Our spatial analysis result were in congruence with our initial Admixture and DAPC 10 clusters partitioning of the GUH panel (**Supplementary Table 14**), exhibiting patterns broadly consistent with isolation by distance where genetic similarity decays with geographic distance (**Supplementary Figure 3a,c)** and corresponds to the DAPC stepping stone model described in Jombart et al (2010).

Our findings in terms of genetic diversity distribution in Brazil is congruent with an earlier report by Alves-Pereira et al. (2018), spatial distribution of genetic diversity of cassava in Brazil is different across bitter and sweet cassava, owing to high haplotype diversity and environmental heterogeneity of cassava growing regions (Alves-Pereira et al. 2018). This is particularly captured in the average cyanide distribution across identified genetic groups (**Supplementary Figure 3f)** in this present study. (Mühlen et al. 2019) and colleagues reported the diverse distributions of sweet and bitter cassava across Brazil, but concluded that the genetic structure of cassava landraces does not coincide with the Brazilian ecogeographic regions (Mühlen et al. 2019). To test (Mühlen et al. 2019)'s hypothesis, we sampled 702 germplasms collected before the year 2000 (excluding breeding lines) and performed spatial analysis to compare the genetic structure of cassava and the ecoregions of Brazil. **Supplementary Figure 11** shows distributions of ancestral coefficients exhibiting similar genetic structure as Brazilian ecoregions while highlighting how migration/germplasm exchanges is modifying the genetic landscape of cassava across the ecoregions of Brazil.

### Genetic diversity estimation

The investigation of MAF distributions across identified groups exhibited variation [suggesting semi-isolated groups] (**Fig. 4a**) and were skewed towards intermediate frequencies (MAF 5–20%) for some groups (SWEET-Group, Amazonas-Group), whereas others (Admix-Group, Bahia-Group, NE-LandRaces, Bitter-Group) show a higher amount of rare (<5%) alleles. The genetic group with the highest diversity (PIC = 0.190) and most number of rare alleles (MAF = 0.144) captured in our dataset is Amazonas-Group, located in the Amazonas ecoregion, consistent with the origin of cassava in Amazonas ecoregion (Allem, n.d.; Clement et al. 2010); as expected from recorded past migration. Diversity decreases away from this epicenter via NE-Admix-Group (PIC = 0.166), in Northeastern region (Coastline of Brazil), to SWEET-Group, BITTER-Group and NE-LandRaces (PIC = 0.166, 0.163 and 0.160), in Northeastern region (covering Cerrado, Caatinga, and Atlantic Forest ecoregions), to WAXY-Group (PIC = 0.159), in Cerrado ecoregion, to Admix-Group (PIC = 0.154), an admixed population covering most of Brazil, possibly resulting from past material migrations along the Amazonas River and along the Atlantic coast to the Southern and Central regions of Brazil. Decreasing numbers of minor alleles are observed along this gradient, suggesting successive small founder populations (stepping stone model).

Overall, MAF provides sufficient power for association studies in a panel that includes a broad range of genetic diversity. SNPs with a MAF of 0.05 (5%) or greater than the average MAF within our population was 0.14 (14%, **Table 1**), highlighting the fact that most loci (SNPs) in this study have required allele frequency (>0.05) for association (and other related trait mapping) studies. The observed MAF highlights informative markers, rare variants and alleles conservation in our population. Our result falls within the range reported for other clonal crops like banana and potato. Sardos and colleagues reported that MAF distribution was skewed toward low frequencies with 31.1% of the markers displaying a MAF below 10% (Sardos et al. 2016). In a more recent study in potato, MAF distribution higher than 10% was also reported (Sharma et al. 2018).

Pairwise *F_ST_* (0.073) reflects a lower population differentiation and a moderate impact of human activities compared to other South American domesticated crops such as tomato (Razifard et al. 2020). The highest level of genetic differentiation (0.108) is observed between SWEET-Group and NE-LandRaces, implying that breeding activities may have restricted gene flow between them and the rest of the groups (**Fig. 4b**). Our results were similar to that of maize, 0.073 (without bottlenecked Bahia-Group) and 0.118 (including bottlenecked Bahia-Group) (Yu et al. 2006). Average *F_ST_* of 0.040 using SSR was reported for cassava germplasm and breeding lines collections of CIAT and African populations (Ferguson et al. 2019). Ferguson (2019), also reported an average *F_ST_* of 0.013 between South American CIAT breeding lines and germplasm collections, indicating a slight differentiation. The overall fixation index *F_ST_*, indicated that the majority of variation is found between groups rather than within groups.

Polymorphism information content (*PIC*) indicates measures of genetic diversity (Botstein et al. 1980; Niu et al. 2019). *PIC* values for this study ranges from 0.142 to 0.375 (with an average of 0.18), in congruence with the maximum *PIC* value of 0.375 for biallelic loci (Nei 1987; Botstein et al. 1980; Luo et al. 2019). Previous studies in cassava reported average *PIC*values of 0.26 (de Oliveira et al. 2014), 0.24 (H. Y. G. de Albuquerque et al. 2018) using SNP markers. In this current study, we only used biallelic SNP, hence the lower *PIC* value estimated when compared to previous studies in cassava. The lower average *PIC* value could also be attributed to low mutation rate in our population (Coates et al. 2009; Eltaher et al. 2018). *PIC* value of 0.177 indicates moderate genetic diversity in our population, given an expected maximum value of 0.375 for markers that are biallelic and 42% of our total SNPs had > 0.2 *PIC* value. Estimate of effective population size, 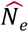 was estimated using an allelic frequency of 0.02 as recommended by Do et al. (2014) in order to increase precision and minimize bias. 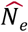 from the identified genetic groups reflects the rate at which variation is lost and may predict the ability to respond to future changes/breeding strategy and significance of conservation approach to genetic diversity and adaptive variation in cassava breeding in Brazil. The estimate of *N_e_* gives a better understanding of the present and future importance of genetic drift of cassava populations and useful information of germplasm management in Brazil. Bitter-Group shows greater genetic diversity and less drift, followed by Admix-Group and NE_LandRaces, while the rest had 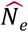 less than 50 and shows more tendencies for genetic drift. Small effective population size of a genetic pool, indicates high LD and potential for improved prediction accuracy (Yabe et al. 2018). Observed heterozygosity was higher for genetic groups having less effective population size (**Table 1**) and this indicates a good measure of the capability of the identified genetic groups to respond to selection, given that the amount of allelic variation left is important for long-term response to selection and survival of populations. In addition, allelic diversity is generally more sensitive to drift than heterozygosity (Allendorf 1986; Sovic et al. 2019), This is made possible in a clonally propagated crop like cassava by incorporated seedlings into clonal stocks to provide new genotypes and additional allelic variation (McKey et al. 2010). We generally observed high heterozygosity across identified genetic groups, indicating a high proportion of genetic variance (allelic variation) across locus in our populations. Taken together, these results show that Admix-Group, Bitter-Group, Amazonas-Group and NE-LandRaces exhibited a higher genetic diversity compared to the rest of the groups.

### Linkage disequilibrium and introgression in Brazilian germplasm

LD in cross-pollinated species decays more rapidly than among self-pollinated species as a result of more effective recombination that occurs in the former (Zhu et al. 2015). The extent of whole genome LD in this study (~107 kb; *r^2^* < 0.1) was higher than previously estimated in Brazilian cassava (~20 kb; *r^2^* < 0.2) (H. Y. G. de Albuquerque et al. 2018), East-West African (~50 kb; *r^2^* < 0.2) datasets (Wolfe et al. 2016) and for HapMap (3 kb; *r^2^* = 0.1) (Ramu et al. 2017) but lower compared to West African (~2 Mb; *r^2^* < 0.1) (Rabbi et al. 2017). Among other outcrossing species, LD observed in this study was higher than LD observed for global panel in maize (~10 kb; *r^2^* < 0.1) (Yan et al. 2009), sweet potato (0.6 kb; *r^2^* < 0.1, 1.2 kb; *r^2^* < 0.2) (Wadl et al. 2018), yam (100 bp; *r^2^* < 0.1) (Akakpo et al. 2017) and potato (~275 bp; *r^2^* < 0.1) (Stich et al. 2013). Given that the magnitude of linkage disequilibrium and its decay with genetic distance determines the resolution of association mapping, chromosomes 5, 13, 17 and 7, 10, 11 show higher and lower LD, respectively, than the average population LD, with chromosome 17 having more than twice the average population LD (**Supplementary Figure 5a, Supplementary Table 7**). The comparison of average LD decay threshold to the standard threshold of 0.1 (*r^2^*) for all genetic pools shows a similar trend with average genetic pool’s LD decay having higher genetic distances across groups (**Supplementary Figure 5b, Supplementary Table 7**). Given the allogamous nature of cassava, the pattern of LD may reduce prediction accuracy, since LD is crucial for genomic selection (GS) breeding (Jannink 2010; Yabe et al. 2018). However, the high LD observed in this study, indicates potential for improved accuracy using this population for GS breeding.

The LD decay, along with the high proportion of SNPs in LD, indicates that association mapping resolution will vary across the genome in accordance with the location of loci associated with the trait of interest within or outside of the region of LD blocks. The observed LD, when compared to earlier studies, could be explained by the additional improvements (introgression) cassava went through in that region (Bredeson et al. 2016; Hahn, Terry, and Leuschner 1980; Okogbenin et al. 2013). In addition, the higher LD when compared to other Brazilian studies, is most likely attributed to the broader population in our dataset, as seen in the presence of extended LD blocks as a result of past introgression effort in some landraces in our population.

Genomic variation across the genome and within groups (excluding family structured group, Bahia-Group) reveals that on average, based on the LD decay (*r^2^* < 0.1), 1,300 to 4,700 SNP markers would be needed to detect large association, while the SNP density of 27,045 was used in this study with 19,085 bp average distance between two SNPs, a much higher marker density will be required to detect small effects association (**Supplementary Table 7**).

The extended LD observed in chromosome 17 (**Supplementary Figure 4**), implying slower LD decay (**Fig. 4d**), was observed across most groups (**Supplementary Figure 6b)**. Excluding chromosome 17 gave genome-wide average LD of 0.019, showing an LD differential of 0.002 from genome-wide average LD (0.021) across all chromosomes, indicating the impact of chromosome 17 to the observed LD decay at genome-wide level. Based on the observed genome-wide LD landscape, we speculated that the extended LD observed in chromosome 17 (**Supplementary Figure 4**), influencing the LD decay for most population groups except group 10 (**Supplementary Figure 6b**), was an introgressed segment. Earlier studies reported that Latin American samples from CIAT showed almost no evidence of *M.glaziovii* introgression on both chromosomes 1 and 4 (Wolfe et al. 2019), where introgressions were detected for African materials (Bredeson et al. 2016; Wolfe et al. 2019). Using the HapMap WGS dataset, we identified “pure” *M. esculenta* ssp*. flabellifolia*, *M. esculenta* ssp. *esculenta* and *M. glaziovii* accessions (Ramu et al. 2017), with *M. esculenta* ssp. *esculenta* accessions made up of Brazilian germplasm. In this study, we identified 294 biallelic ancestry-informative single-nucleotide markers that represent fixed (or nearly fixed) differences between *M. esculenta* ssp. *esculenta* and *M. esculenta spp. flabellifolia* (**Supplementary Figure 8a)**. However, the identified variants were too few and too sparse to assign segments of *M. esculenta* ssp*. flabellifolia* ancestry across the genome. On the other hand, when *M. glaziovii* was compared to *M. esculenta* Brazilian germplasm, we identified a large set of differentiating markers, in contrast with the few differentiating markers earlier reported for CIAT germplasm (Wolfe et al. 2019). The observed 3,238 intersecting SNPs in our GBS dataset may indicate evidence of past or more recent introgression in Brazilian germplasm (**Supplementary Figure 8b-c)**. Bredeson and colleagues (2016), hypothesized that some of the 98 described species of *Manihot* (O. K. Miller and Singer 1972; Nassar, Hashimoto, and Fernandes 2008) may represent interspecific hybrids or admixtures (Bredeson et al. 2016). This hypothesis was further developed in (Ramu et al. 2017). In the current study, we found individuals that are interspecific hybrids of *M. glaziovii* × *M. esculenta* (“MEPRUNOSA351E058” and “EPRUNSA351xMGLAZIOVII”), and another individual (“UFB_002”) that appears to be from a “pure” *M. esculenta* × *M. glaziovii* hybrid backcrossed to another *M. glaziovii* (or, alternatively, a hybrid that includes *M. esculenta* that already had introgression), supporting the hypothesis put forward by Bredeson et al. (2016). “UFB_002” and a couple of others (“M1_18:250492593” and “ManipebaPreta253:250437596”) share alleles with neither of *M. esculenta* or *M. glaziovii*, meaning, they may be (hybrids with) a third species. In addition, Bredeson and colleagues (2016), found an unidentified ancestral segment shared by two Brazilian germplasms that could not confidently assign their ancestry based just on the collection of *M. esculenta* and *M. glaziovii* alleles used in their study. We found evidence for introgression of *M. glaziovii* in Brazilian Germplasm collections on chromosome 17 and genetic Bitter-Group (**Fig. 5**), confirming our initial speculation of introgression on chromosome 17 based on LD landscape. This might be due to one or more differences in chromosome structure (such as an segmental inversion) between *M. glaziovii* and *M. esculenta* (preventing viable offspring with recombinant homologs) that is/are unobservable using only short Illumina resequencing reads. The individuals in Bitter-Group, represent landraces of bitter cassava (see **Supplementary 3f** for average HCN per group) from Northeastern region of Brazil, indicating that the introgression segment we observed may have been as a result of past breeding effort rather than a more recent breeding activity (Allem, n.d.). Wolfe and colleagues (2019), reportedly found a segment of *M. glaziovii* in three CIAT accessions originating from Brazil (“BRA134:250327456”, “BRA172:250327458” and “BRA842:250327480”). We could confirm this finding, these accessions clustering within Bitter-Group. Explaining the ancestral segments in **Fig. 5,** with blue and green segments highlighting the presence of *M. glaziovii* introgressions, specifically in individuals identified as hybrid including “Manipeba”, clones known rustic form of cassava from Northeast Brazil, hypothesized as a transitional link between wild ancestor and cultivated cassava (Allem, n.d.; M. de Albuquerque 1969). The high 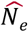 observed in Bitter-Group may be inflated by such introgression.

### Genome Scan for Selection

Substantial natural and artificial selection was revealed by the genome scan for selection, highlighting genetic variation of cassava collection in Brazil. The genome-wide scan for selection based on population differentiation in the nine ancestral populations detected (from spatial analysis), showed that the QQ-plot provided evidence that confounding errors were accounted for. The Manhattan plot exhibited islands of strong differentiation around positions in all chromosomes (except chromosomes 3, 6, 8, 10, 11, 14 and 17), with 24 differentiating loci identified across the genome (**Supplementary Figure 12**).

Crop domestication in most cases involves intermediates that are unexplored or underexplored corresponding to historic evolutionary stages in domestication events, and cassava is not an exception. The southwestern Amazonia indicates region around the center of domestication of cassava as was earlier hypothesized and supported by (K. M. Olsen and Schaal 1999; K. Olsen and Schaal 2001; Kenneth M. Olsen 2004; Mühlen et al. 2019). The robustness of the inferred nodes in **Fig. 7** and the fraction of the explained variance for three migration events, statistically supports the identified population splits and gene flow events. (Walker et al. 2019), showed the Amazonas ecoregion, the source of the rivers that feeds its rivers and its connections to other ecoregions within Brazil**. S**uggesting the likely connections between the Amazon ecoregion and the Amazon basin, leading to potential route of seeds/cuttings migration or dispersion by animals or other elements. Given that Northern Bolivia and Amazonian (Clement et al. 2010) shows ancient tracks of cassava domestication, supporting the observed gene flow events, with migration through the Amazonas to the rest of the ecoregions potentially via the river that connects to the Amazon basin. This tree seeks to suggest a second population split between Caatinga and Atlantic forest. However, Caatinga seems to be slightly earlier than Atlantic forest based on the genetic drift parameter. Leading us to suggest that cassava may have been propagated to the coastal part of Brazil independently of its migration through the Amazon region. This hypothesis is supported by an independent spatial analysis of this set of individuals, with Southern Amazonas showing the same ancestral coefficient as those in the coastal region of Brazil (**Supplementary Figure 11b**). Atlantic forest ecoregion may have experienced less genetic drift given that breeding activities had mostly occurred around that region in the last four decades. The gene flow from Amazonas to Cerrado and Atlantic forest is consistent with Olsen and Schaal 1999. We don’t have sufficient information to make inference on the third gene flow from Pampa to Atlantic forest ecoregion.

## Conclusion

In this study, we characterized genetic diversity and investigated population structure, duplicate analysis, linkage disequilibrium, population differentiation and estimate of effective population size among the identified genetics groups in Brazilian cassava. Accurate identification of crop cultivars is crucial in assessing the impact of crop improvement program outputs, and highlighting the magnitude of genetic drift amongs identified genetic pools and its impact on survival, reproduction, and the rate at which a population enters the extinction vortex. Applying the 50/500 rule as described by (Franklin and Frankham 1998), genetic group’s effective population size of at least 50 individuals is necessary for conservation of genetic diversity in the short term and avoids inbreeding depression. While a genetic group’s effective population size of at least 500 individuals is required for long term survival with the ability to evolve under a changing environment and avoiding serious genetic drift. Minimizing loss of genetic variation via genetic drift should be an important objective of the cassava breeding program in Brazil. This can be partly achieved by minimizing inbreeding as a result of equalisation of family sizes in germplasm collections and regeneration. Based on our datasets, we suggest that: (1) there are 1,536 IBS selected unique accessions that represents the diversity within Brazilian germplasm collection. (2) there are 5 major ancestral populations and 10 genetic groups in the Brazilian germplasm collections. (3) selection based on traits (ie: waxy, sugary and bitter related traits) of interest by breeders and farmers could have led to the observed population structure in the collected germplasm across Brazil. (4) Spatial analysis showed with consistency the Southwestern Amazonia as the center of domestication of cassava, highlighting the cassava diversity landscape of Brazil, including impact of breeding activities. (5) LD decay across the genome indicates that interval of 1,300 to 4,700 SNP markers are required to detect with reasonable power, association of large quantitative trait loci (QTL) effect, based on our GBS dataset. (6) In addition, this defines the SNP density for development of PCR amplicon sequencing for genotyping and also suggest that GBS can be done at higher DNA multiplexing, thereby reducing the costs of variety fingerprinting. (7) We found evidence for *M. glaziovii* introgression in Brazilian germplasm which may have resulted from past breeding activities. (8) The identification of *M. glaziovii* introgression (and a possible third species sampled by UFB_002) in Brazil means maybe that interspecific hybridization is a more wide-spread phenomenon than previously suspected, and an effort to sequence and assemble the complete genomes of related *Manihot* species would enable a more comprehensive survey for sources of desirable alleles/traits. (9) We suggest that cassava may have been propagated to the coastal region of Brazil independently of its migration from the Amazonas ecoregion. Several past studies identified the origin of cassava domestication to be the Southwestern Amazonia, our findings provide the first novel evidence of cassava domestication using spatial analysis to project the ancestral coefficient on Brazilian map and further show the evidence of divergence of an ancestral population into two ancestral groups.

Studies with domesticated animals and plants have shown that the loss of genetic variation has negative effects on growth, survival, development and also reduces the evolutionary potential of a population (Allendorf 1986; Greenbaum et al. 2014). This is also important for germplasm collections that serve as a gene bank for possible reintroduction into the breeding program. In this study, we measured genetic variation and estimated effective population size across identified genetic groups to inform optimized germplasm management and conservation of genetic variability for prospecting of breeding approaches in a cassava breeding program. Breeding schemes should be designed to increase effective population size, thereby minimizing genetic drift in cassava germplasm in Brazil. Genetic variability fuels breeding innovation. In an ever-increasing globalization context, public and private breeding initiatives foster germplasm exchange to solve key breeding challenges (ie: disease resistance). With Brazil as a center of diversity and Africa as the main production zone, the cassava breeding community illustrates well this challenge. Such action implies a better characterization and tracking of cassava germplasm including reassignment of new names to exchanged germplasm in breeding programs.

In a context where Amazonia’s *In situ* genetic resources are facing increasing conservation challenges, present findings present significant progress in characterizing *Ex situ* cassava germplasm collection. Results including identification of crop cultivars for efficient germplasm management, SNP density requirement for the development of simple PCR amplicon assay for genotyping, and provision of important information on the genome-wide assessments of the genetic landscape of cassava in Brazil as future resources for breeding activities; such as genomic selection, association studies and marker-assisted selection to increase genetic gain in cassava breeding programs in Brazil.

## Availability of data and material

Genotyping (SNP) data used in this study for 3,345 accessions were deposited on cassavabase.org hosted at “ftp://ftp.cassavabase.org/manuscripts/Ogbonna_et_al_2020/population_structure_manuscript”.

## Author contribution statement

Designed experiment: AO, EO, GB; Performed experiment: AO, GB, LBRA. Project supervision: LM, GB, EO; First draft of the manuscript: AO; All authors reviewed and approved the manuscript.

## Acknowledgements

The authors appreciate Jean-Luc Jannink and Deniz Akdemir both from Cornell University, Ithaca, NY for providing helpful feedback. The authors thank Jessen V. Bredeson from University of California, Berkeley, for providing helpful discussion and feedback.

## List of tables and figures

**Supplementary Table 1:** panel definitions. GBS All (GA), GBS unique (GU), GBS All & HapMap (GAH), GBS Unique and HapMap (GUH), GBS All & HapMap and GPS coordinates (GAHg).

**Supplementary Table 2:** 120 randomly sampled individuals from identified groups.

**Supplementary Table 3:** List of HapMap clones *M*. *esculenta*, *M*. *esculenta* ssp*. flabellifolia*, *M*. glaziovii used for introgression detection.

**Supplementary Table 4:** List of germplasm across the ecoregions of Brazil used in population split analysis.

**Supplementary Table 5:** List of non-duplicated selected for GBS-Unique (GU) panel.

**Supplementary Table 6:** GBS-all panel (GA & GAH) and GBS-unique panel (GU & GUH) metadata.

**Supplementary Table 7:** Linkage disequilibrium and physical map distance for which trend lines of non-linear regression falls below R = 0.1 and average R value across the genome for each genetic pool.

**Supplementary Table 8:** list of biallelic ancestry-informative single-nucleotide markers that represent fixed or nearly fixed differences between *M. esculenta* ssp. *esculenta* and *M. esculenta* ssp*. flabellifolia*.

**Supplementary Table 9:** list of biallelic ancestry-informative single-nucleotide markers that represent fixed or nearly fixed differences between *M. esculenta* and *M. glaziovii*.

**Supplementary Table 10:** List of *M*. glaziovii ancestry-informative markers from HapMap (Ramu, 2017) intersecting with 27,094 SNP GBS dataset.

**Supplementary Table 11:** List of individuals and their corresponding biomes.

**Supplementary Table 12:** List of 702 individuals excluding breeding lines used in Spatial population structure analysis.

**Supplementary Table 13:** *F*_*st*_ genome-wide scan loci above the bonferroni threshold of 5.732715.

**Supplementary Table 14:** DAPC groups for the GBS Unique-HapMap (GUH) panel for unique individuals used in GUHg panel.

**Supplementary Figure 1:** phylogeny for cassava clones included in the GU panel.

**Supplementary Figure 2:** Plot of principal component from DAPC analysis.

**Supplementary Figure 3:** Inference of the number of clusters in GUH panels.

**Supplementary Figure 4:** Local pattern of linkage disequilibrium.

**Supplementary Figure 5:** Average linkage disequilibrium and LD-decay

**Supplementary Figure 6:** Trend line of the non-linear regression of the linkage disequilibrium measure.

**Supplementary Figure 7:** Genetic group 2 Identity-by-Descent estimation.

**Supplementary Figure 8:** Distribution of bi-allelic ancestry-informative single-nucleotide markers.

**Supplementary Figure 9:** Inferred segmental ancestry showing introgression.

**Supplementary Figure 10:** Spatial analysis and ancestral coefficients.

**Supplementary Figure 11:** Spatial Analysis of 702 germplasm collected before the year 2000.

**Supplementary Figure 12:** Genome-wide scan for selection.

**Supplementary Note 1:** Bahia-Group discussion.

