## Supplementary Figures for "Comprehensive genotyping of Brazilian Cassava (*Manihot esculenta* Crantz) Germplasm Bank: insights into diversification and domestication"

The following Supporting Information is available for this article:

**Supplementary Figure 1:**

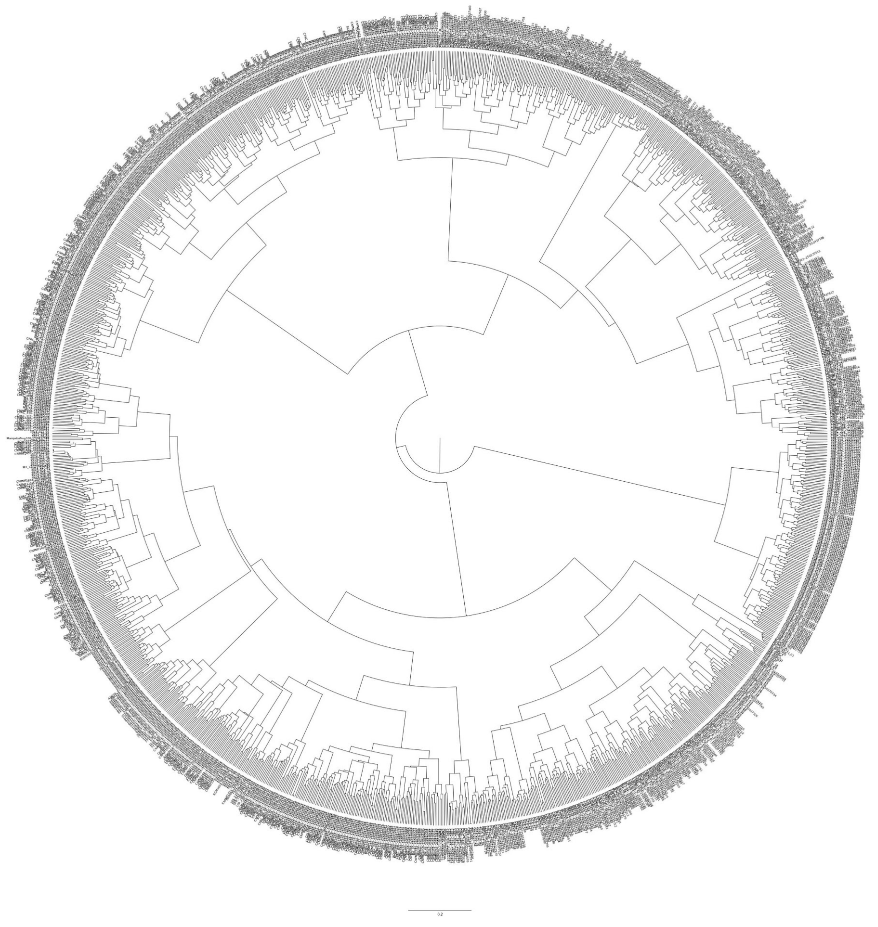

Phylogeny for cassava clones in the GU panel, the IBS-selected unique core set made of 1,536 individuals. The phylogeny tree was constructed from Identity-by-State (IBS) pairwise distance using hierarchical clustering hclust function ("ward.D2" method) in phyclust v0.1-28 R package version 3.6.3 (2020-02-29) and visualized in FigTree v1.4.4 (http://tree.bio.ed.ac.uk/software/figtree/).

**Supplementary Figure 2:**

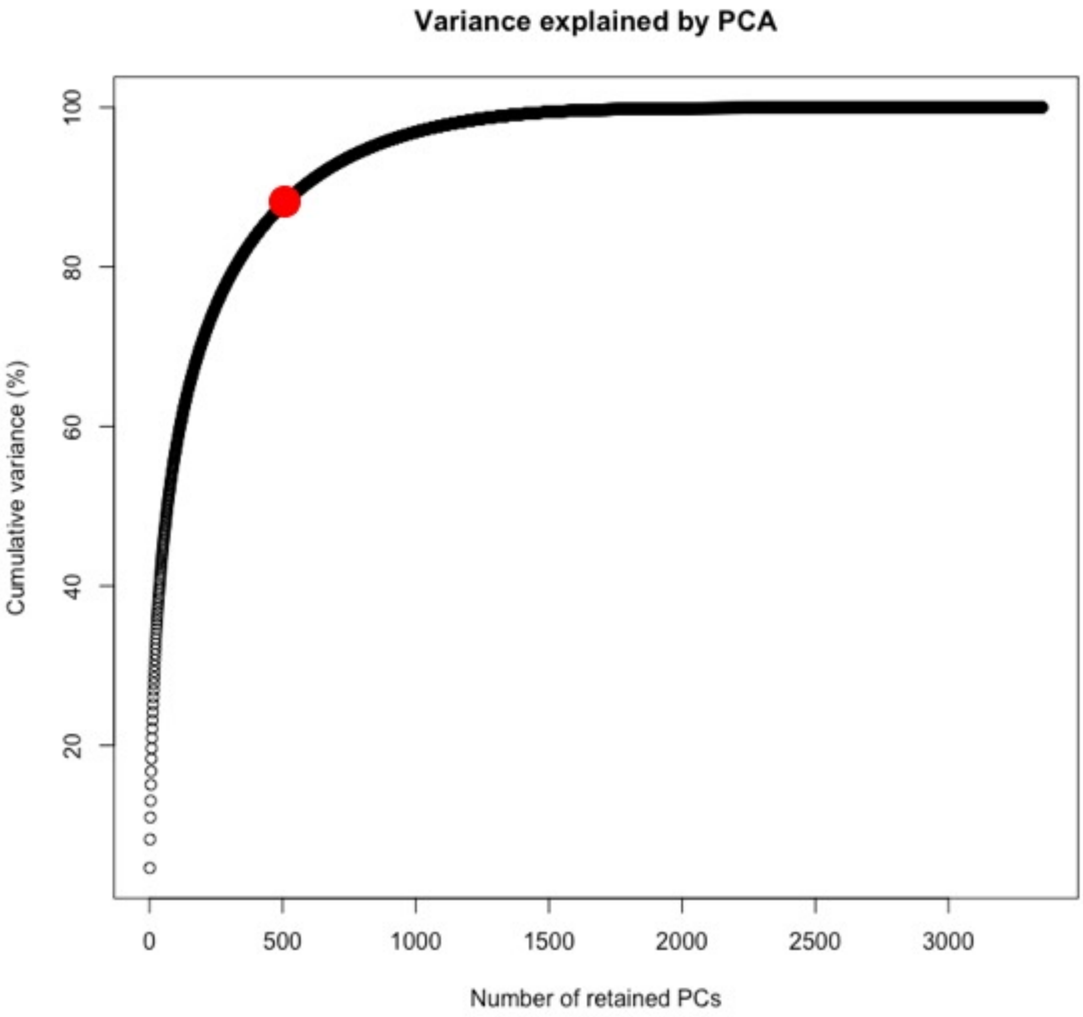


Plot of principal components from DAPC analysis. 500 PCs were used explaining about 90% genetic variation in our dataset. 500 PCs were retained. The red dot indicates the number of PCs retained and the equivalent variance explained.

**Supplementary Figure 3:**

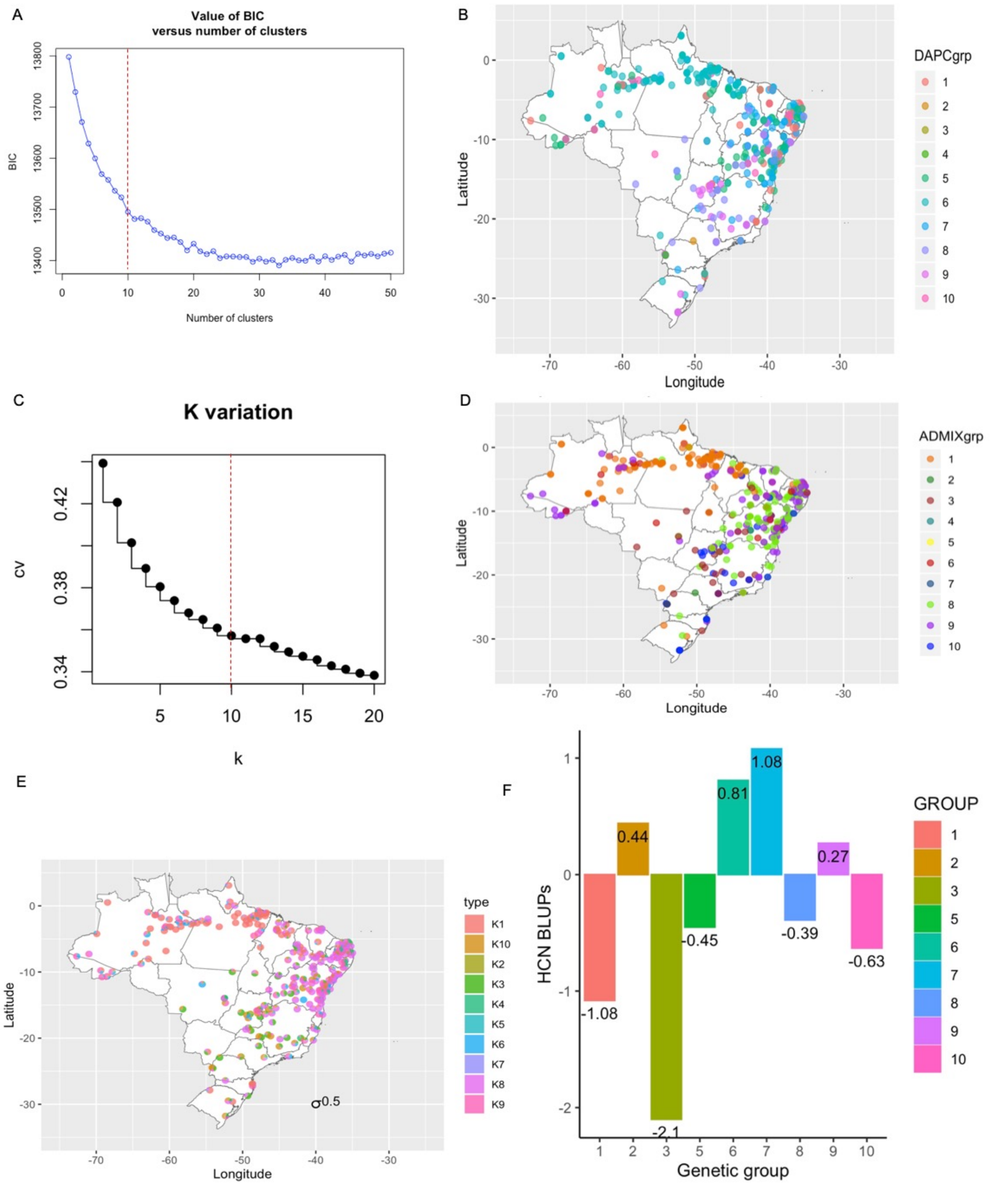


Discriminant Analysis of Principal component (DAPC) and Admixture for GUH panels. Group distributions were plotted using 1,039 accessions having global positioning systems (GPS) in the GUH panel. (**A**) Discriminant analysis of Principal Components (DAPC) Bayesian information criterion (BIC) plot. Ten clusters were chosen for DAPC based on the Bayesian information criterion (BIC) and indicated by the dotted red line. (B) Genetic group distribution for GUH panel for 1,039 individuals. DAPC analysis was performed on the complete panel, while group distribution was accessed using only individuals with GPS coordinates. (C) Admixture 5 folds cross-validation error plot for change in K. K = 10 clusters were selected for Admixture based on the standard error of the 5-fold cross validation. The actual number of populations (K) is indicated by the dotted red line. (D) Proportion of admixture for GUH panel for 1,039 individuals with GPS coordinate. Admixture genetic groups geographical distribution and proportion of admixture across each genetic group for GUH panels for 1,039 individuals with GPS coordinates. Admixture analysis was performed on the complete panel while group distribution was accessed using only individuals with GPS coordinates. (E) Proportion of admixture for GUH panel for 1,039 individuals with GPS coordinate. Admixture proportion is displayed on Brazilian map using a pie chart of 0.5 radius for each individual in our panel with GPS coordinate. The legend key K1 through K10, represents the different ancestral coefficient. (F) Barplot of average hydrogen cyanide content per identified genetic group. Plotted on Y-axis are the Best Linear Unbiased Predictions of HCN and on the X-axis are the identified genetic groups. HCN values for individuals in the genetic groups are accessed from (Ogbonna et al. ).

**Supplementary Figure 4:**

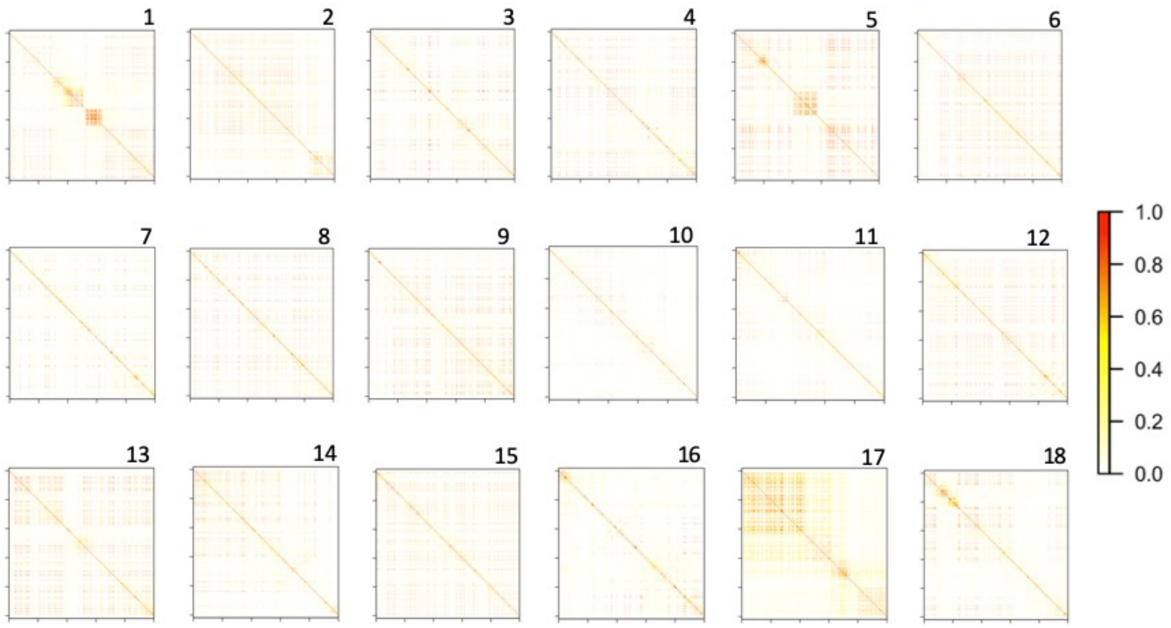


Local pattern of linkage disequilibrium ($r^2$) along each of the 18 cassava chromosomes, showing genome landscape of Brazilian germplasm. Note the large linkage disequilibrium blocks in chromosomes 1, 5, and 17. Single nucleotide polymorphisms are arranged according to their order and not their physical position or genetic distance.

**Supplementary Figure 5:**

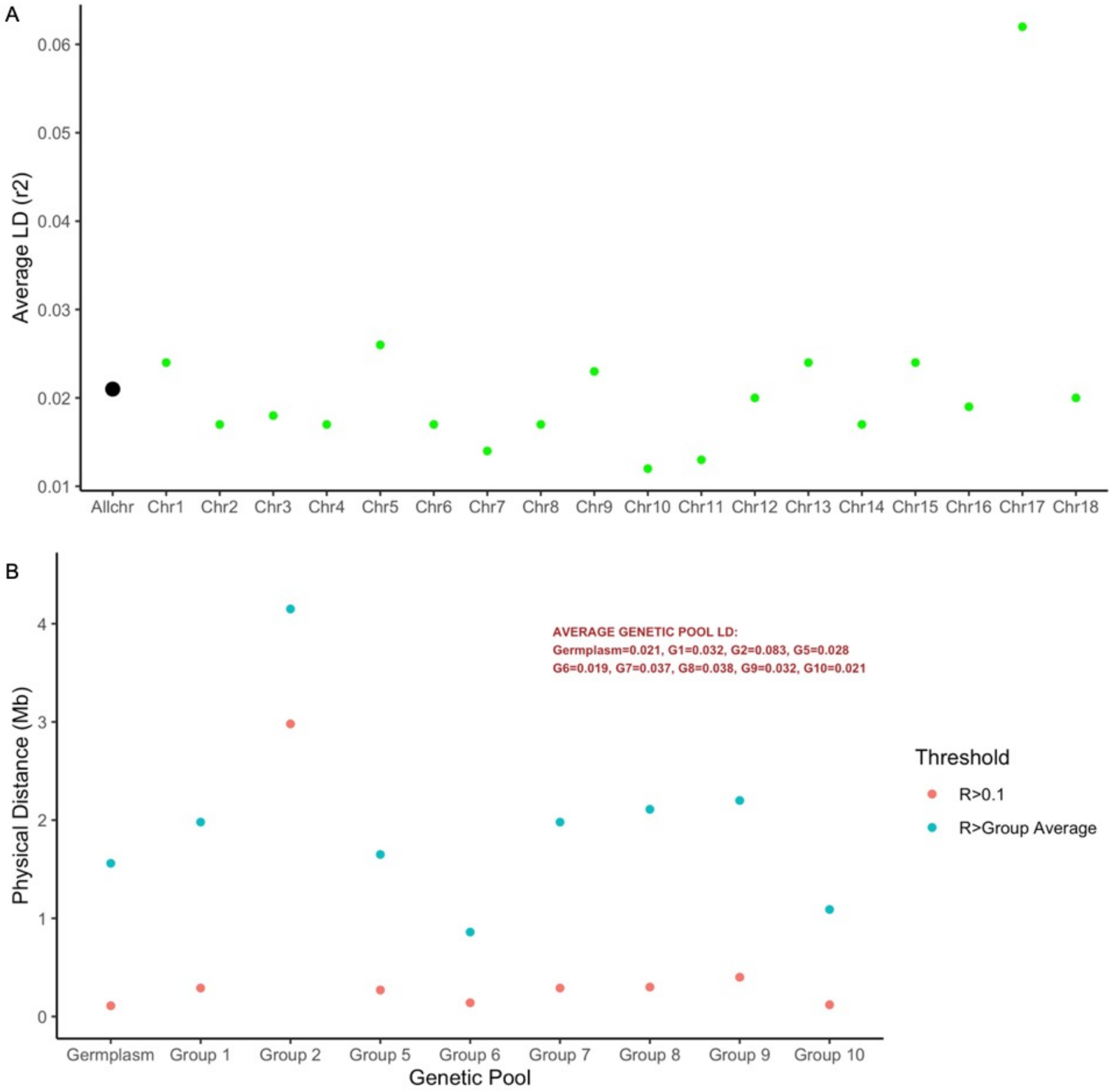


Average linkage disequilibrium and LD-decay of Brazilian germplasm and genetic groups based on Hill and Weir model (*$r^2$*). (A) Average Linkage disequilibrium for the whole genome (black dot) and all chromosomes (green dot). The average LD was computed using Group 1,5 - 10. Family-structured-group 2 was excluded along with group 3 and 4 with <100 individuals. (B) Physical distance (LD decay) for which the trend line of the non-linear regression falls below $R^2$ = average Brazilian germplasm, or below $R^2$ = 0.1 for Brazilian germplasm and each identified genetic group.

**Supplementary Figure 6:**

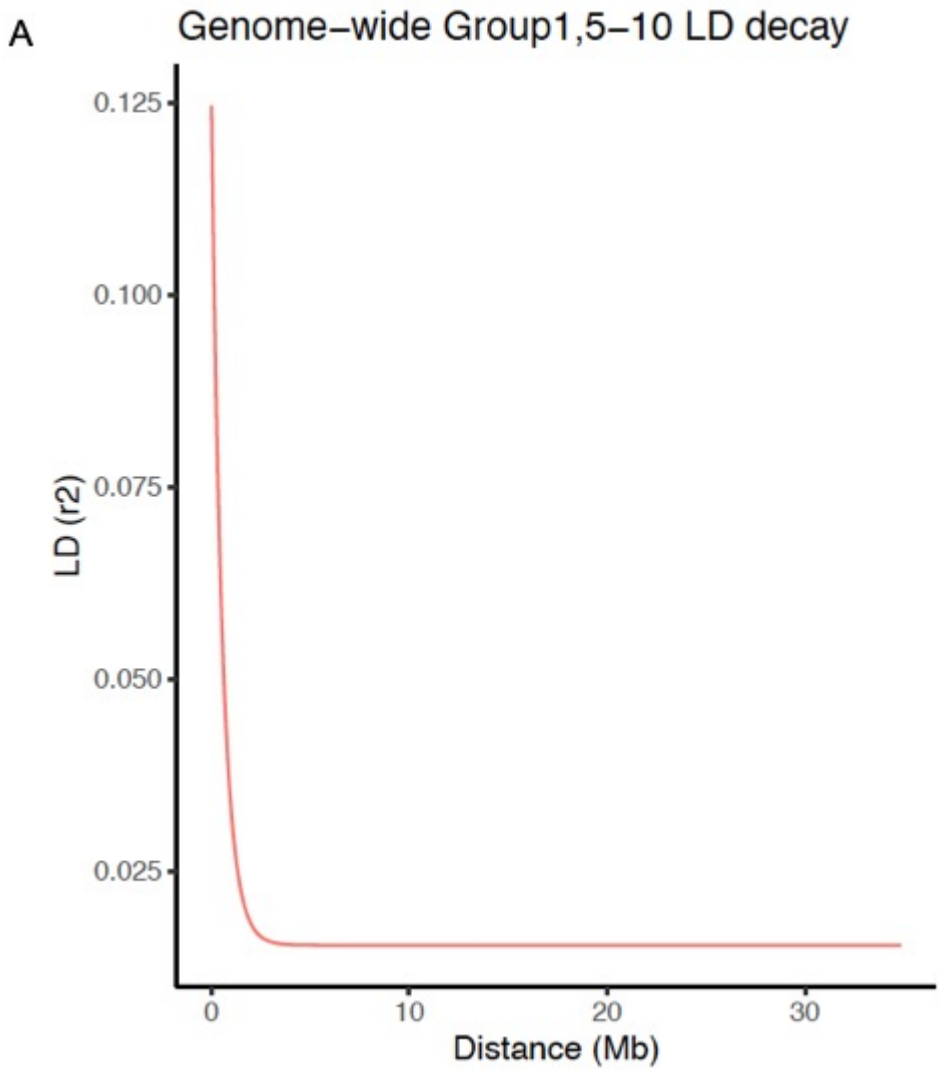

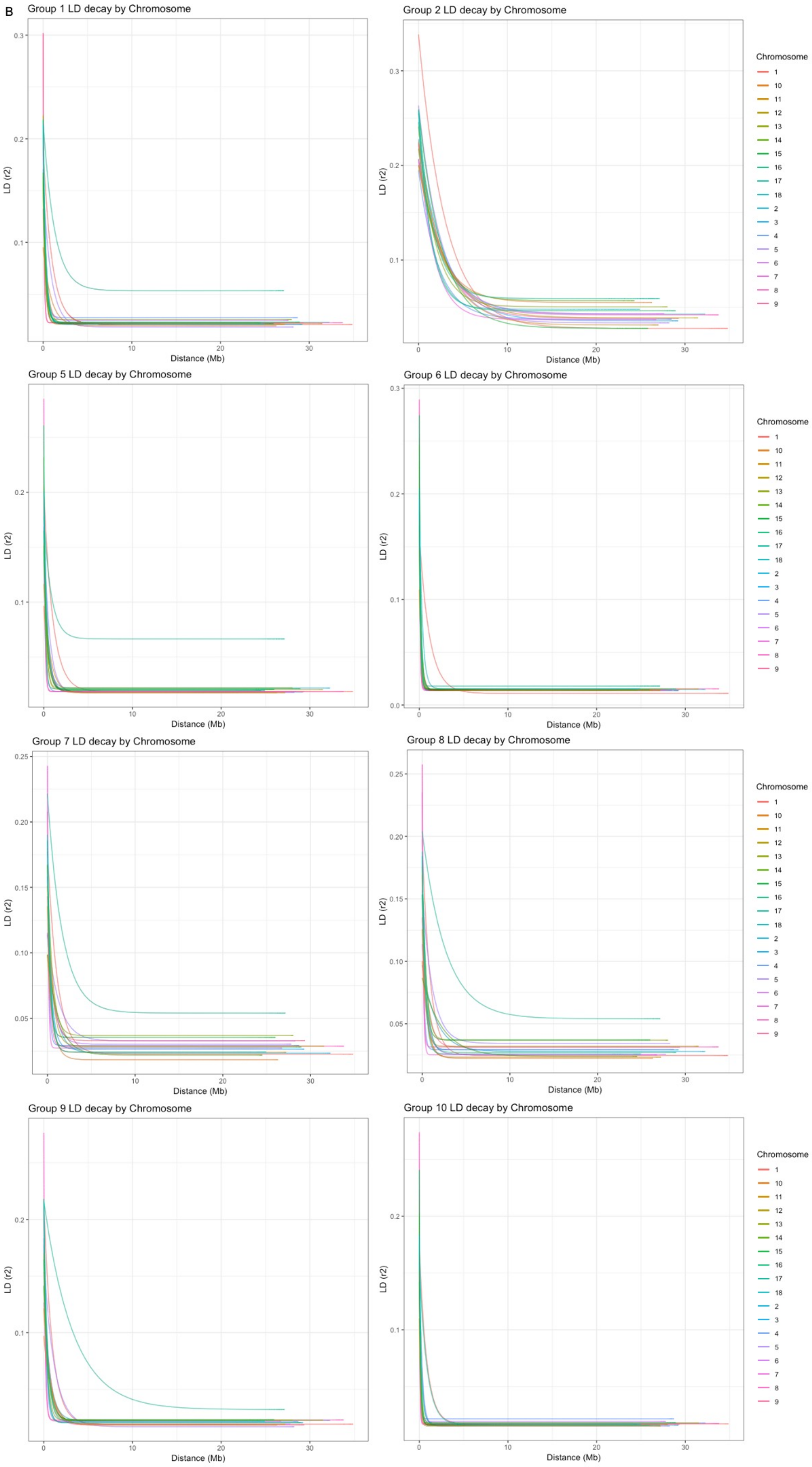
B
Group 1 LD decay by Chromosome
Group 2 LD decay by Chromosome
Group 5 LD decay by Chromosome
Group 6 LD decay by Chromosome
Group 7 LD decay by Chromosome
Group 8 LD decay by Chromosome
Group 9 LD decay by Chromosome
Group 10 LD decay by Chromosome
LD (r2)
Distance (Mb)
Chromosome
1
10
11
12
13
14
15
16
17
18
2
3
4
5
6
7
8
9

(A-B) Trend line of the non-linear regression of the linkage disequilibrium measure $r^2$ versus physical distance (Mb) between single nucleotide polymorphism (SNP) marker pairs across the genome and the chromosomes for each identified subgroup in the genome of the Brazilian germplasm. The LD is computed chromosome-wide. The average LD $r^2$ is 0.021 and drops to background level ($r^2 < 0.1$) at around 107 kb across the genome.

### Supplementary Note 1:

Bahia-Group (group 2) showed a clear departure from other groups (based on LD-decay, MAF and Fst- **Fig. 4a-c**). To further explore the family structure observed and genetic ancestry between individuals in group 2, **Supplementary Figure 7** shows within family relationship heatmap for genetic group 2 with an average estimated kinship coefficient of 0.4 (min=0, max=0.7), highlighting shared haplotypes between individuals initially originating from a biparental cross. Using the threshold of 0.45 or greater as described by (Myles et al. 2011; Bredeson et al. 2016) to define individuals sharing putative first-degree relationships, 34% of individuals in group 2 have first-degree relationships (**Supplementary Figure 7b-c**). 100% of the individuals showed parent-offspring relationships with expected probabilities of IBD0, IBD1, and IBD2 of 0.0, 1.0, and 0.0, respectively. No individual showed a full-sibling relationship, given the expected probabilities of IBD0, IBD1, and IBD2 of 0.25, 0.5, and 0.25, respectively.

**Supplementary Figure 7** confirmed the family structure observed in group 2 by highlighting the few individuals that were involved in the development of the population. The self-reported parent-offspring pairs show that they shared a reasonable amount of their alleles IBD=1, from completely related to relatively closely related, hence clustering mostly around the top-right of the **Supplementary Figure 7b**. At the bottom right of the plot where Pr(IBD=0)=1 and Pr(IBD=1)=0, indicated that the two unrelated individual does not share any alleles identical by descent at every locus across the genome and represented the initial parents of the biparental cross that was done for the progenies that were further used in developing this group. This same information was also captured in the first two individuals on the left of the heatmap (**Supplementary Figure 7a)**.

### Supplementary Figure 7:

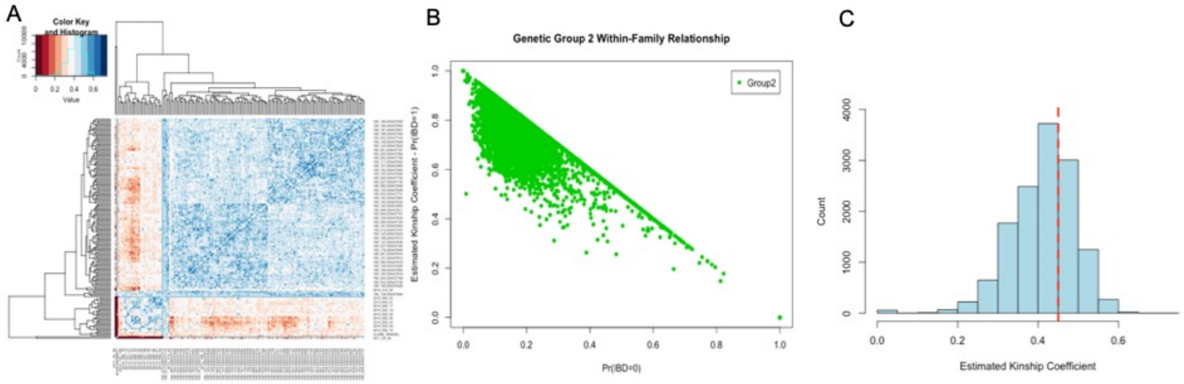


Bahia-Group (Genetic group 2) Identity-by-Descent estimation used for assessing genetic homogeneity across all pairwise comparisons of individuals. **(A)** Heatmap of PI_HAT values across pairwise of all individuals in group 2. (B) Plot of having probability IBD of 0 on the X-axis by probability of having IBD of 1 on the Y-axis (estimated kinship coefficient). **(C)** Histogram of estimated kinship coefficients ($\pi$) for all pairwise relatedness comparisons between the group 2 accessions. The red vertical broken line indicates the threshold of 0.45 for first-degree relationship between individuals. We inferred within-group homogeneity across all pairwise combinations of sampled individuals in genetic subgroup 2 using the IBD coefficient PI_HAT implemented in PLINK v.1.9.

**Supplementary Figure 8:**

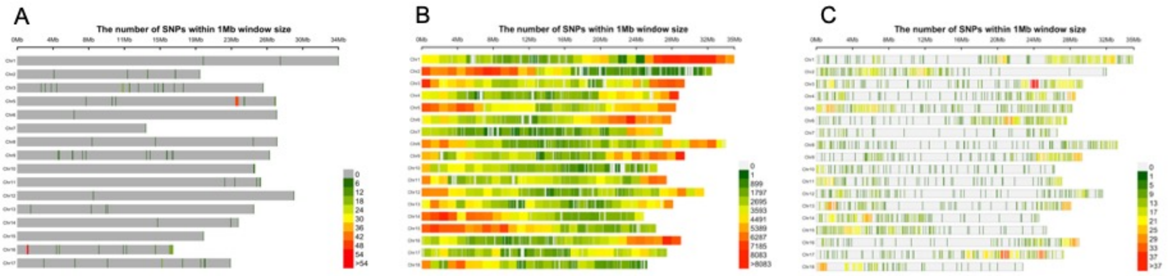


Distribution plots of bi-allelic ancestry-informative single-nucleotide markers that represent fixed, or nearly fixed, differences between individuals of two different species of cassava. (A) The plot shows the distribution of 294 marker differences between *M. esculenta* and *M. flabellifolia* in the HapMap WGS dataset. (B) The plot shows the distribution of 1,795,898 marker differences between *M. esculenta* and *M. glazovii* in the HapMap II WGS dataset. (C) The plot shows the distribution of 3,238 markers different between *M. esculenta* and *M. glazovii* resulting from the intersection of our GBS and WGS HapMap II datasets. The legend scale indicates the number of SNPs within 1Mb window size.

**Supplementary Figure 9:**

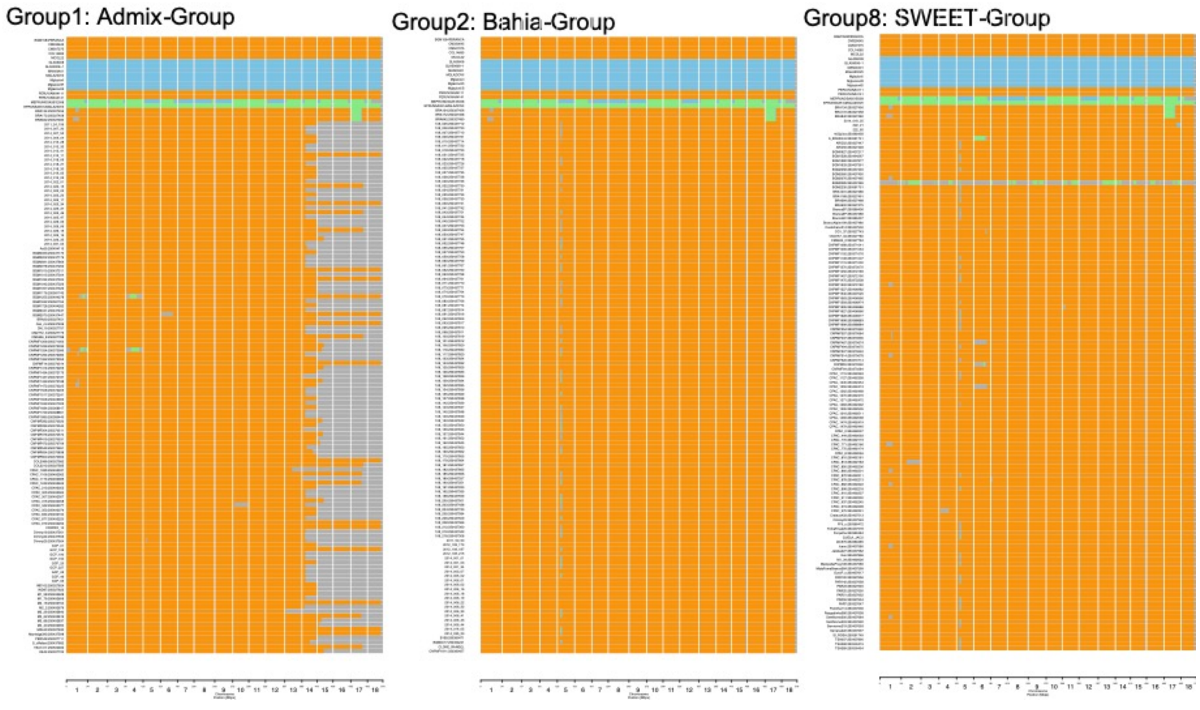

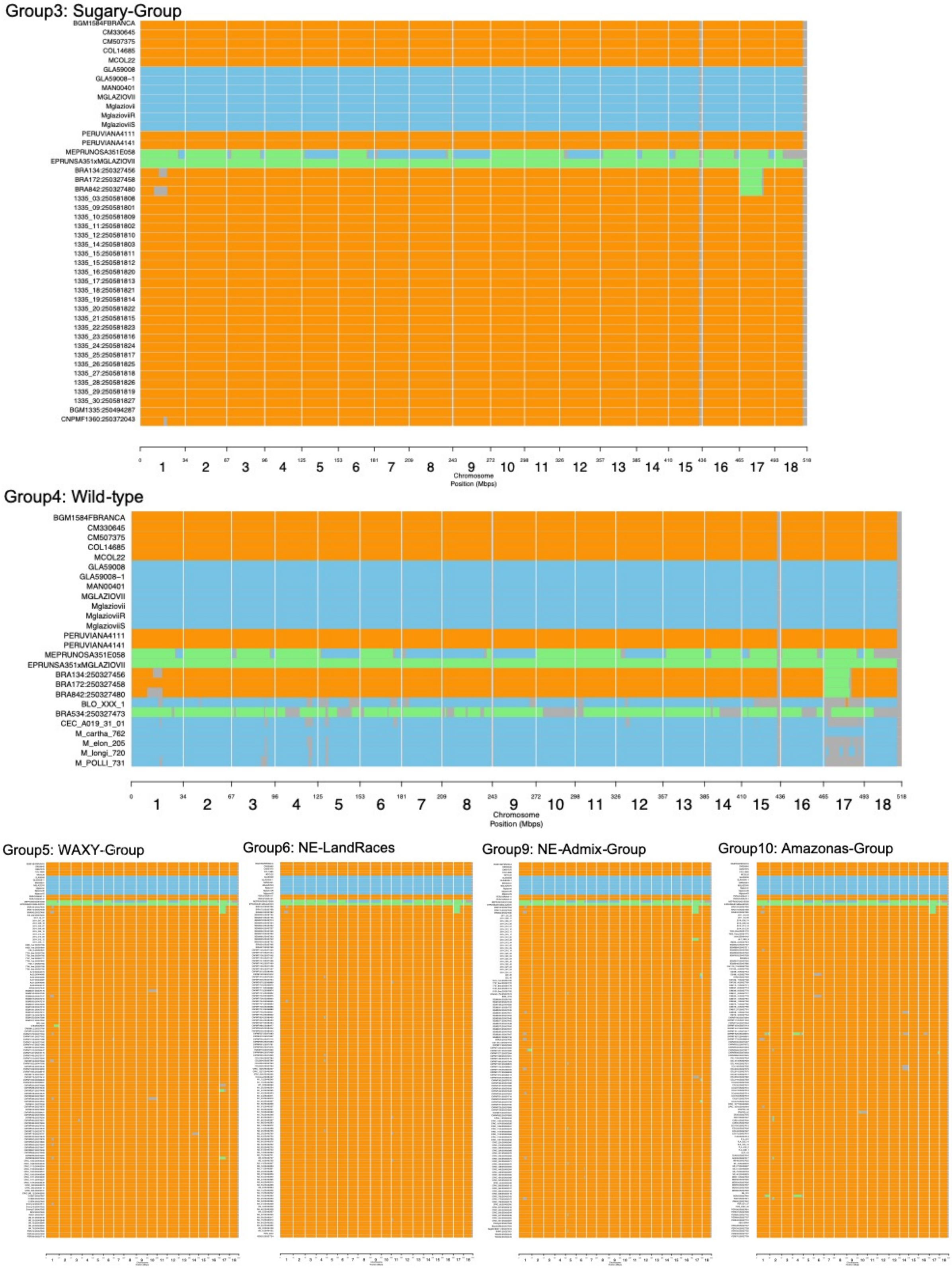


Inferred segmental ancestry showing introgression of M. glaziovii in chromosome 5 inferred from GBS dataset. Introgression plot of Bahia-Group and SWEET-Group (Group 2 and 8) shows introgression of M. glaziovii in chromosome 5. Genetic group 2 are made up of a family structured population, while genetic group 8 are made up of sweet cassava accessions. The orange color code means both haplotypes are M.*esculenta*; blue is both haplotypes M.*glaziovii*; green is one haplotype *M. esculenta*, one *M.glaziovii*; grey means "no call" or "no information". The rest of the groups had no introgression segment.

**Supplementary Figure 10:**

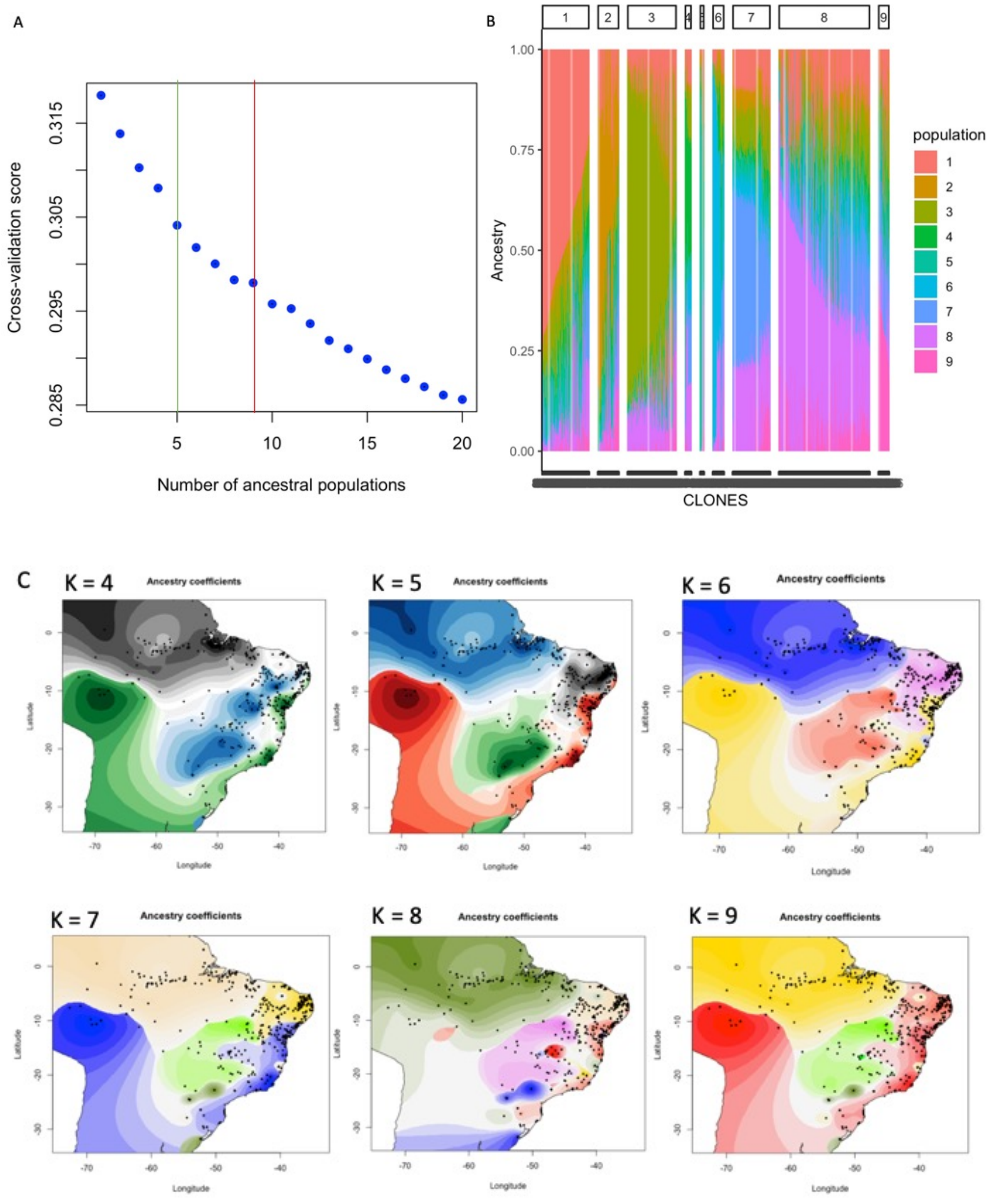


Analysis of 1,657 accessions using tess3r. (A) Cross validation error as function of number of ancestral populations, K. The green and red lines show the number of major and total ancestral populations K = 4 and 9, respectively. (B) Barplots of ancestry coefficients with K = 9 ancestral populations. (C) Geographic maps of ancestry coefficients for K = 4 - 9 ancestral populations.

**Supplementary Figure 11:**

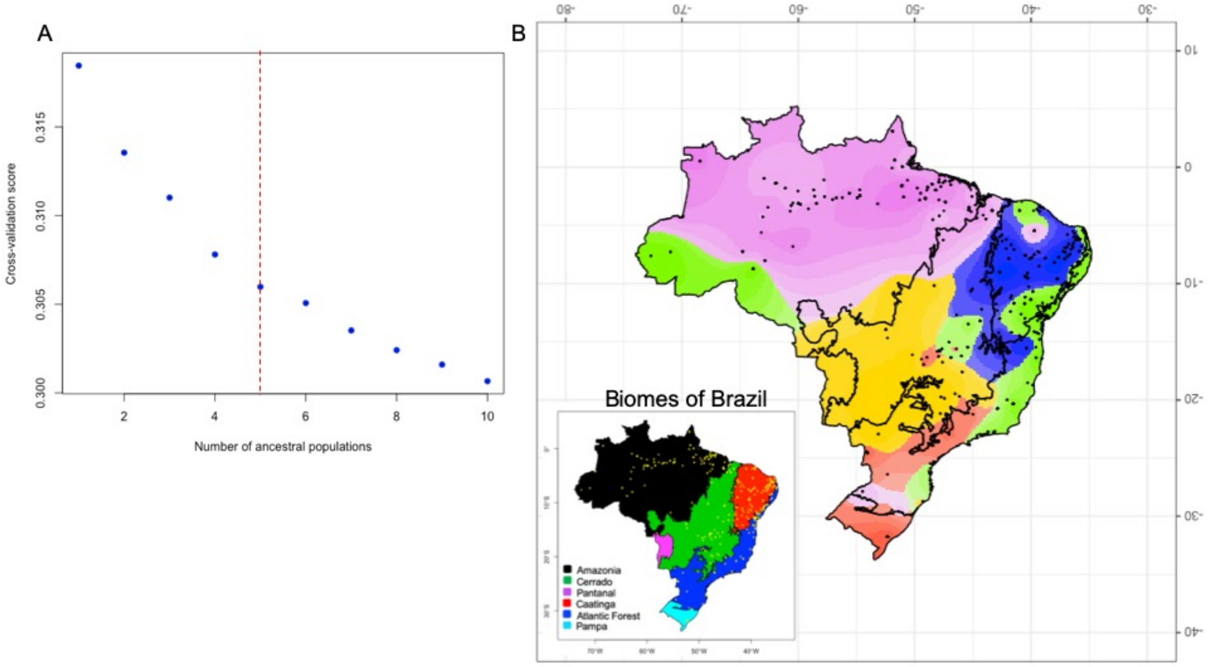


Spatial Analysis of 702 germplasm collected before the year 2000. **(A)** Cross validation error as function of number of ancestral populations, K. The red lines show the number of major total ancestral populations K = 5. **(B)** Geographic maps of ancestry coefficients for K = 5 ancestral populations. Bottom left ecoregions (biomes) of Brazil are indicated.

**Supplementary Figure 12:**

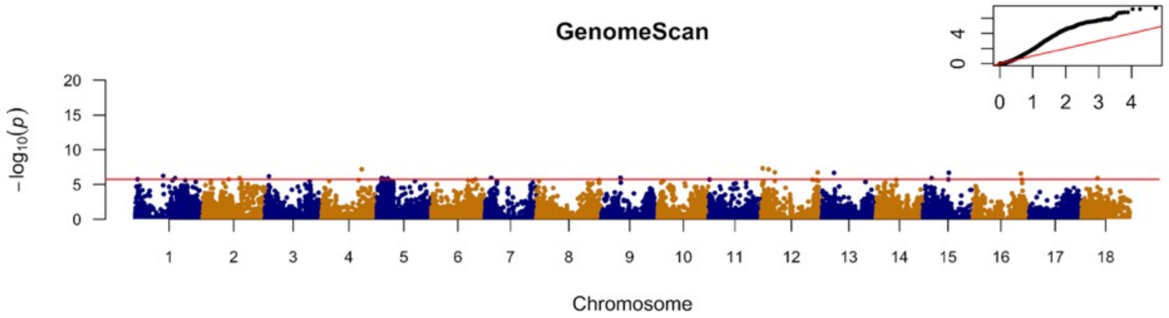


Manhattan plot of log10(P-values) for genome-wide scan for selection on 1,657 accessions using tess3r. At K = 9, 24 significant SNPs were detected to be above Bonferroni threshold (-log10(0.05/27020) = 5.732715).
